# Heritable microbiome variation is correlated with source environment in locally adapted maize varieties

**DOI:** 10.1101/2023.01.10.523403

**Authors:** Xiaoming He, Danning Wang, Yong Jiang, Meng Li, Manuel Delgado-Baquerizo, Chloee McLaughlin, Caroline Marcon, Li Guo, Marcel Baer, Yudelsy A.T. Moya, Nicolaus von Wirén, Marion Deichmann, Gabriel Schaaf, Hans-Peter Piepho, Zhikai Yang, Jinliang Yang, Bunlong Yim, Kornelia Smalla, Sofie Goormachtig, Franciska T. de Vries, Hubert Hüging, Ruairidh J. H. Sawers, Jochen C. Reif, Frank Hochholdinger, Xinping Chen, Peng Yu

## Abstract

Beneficial interactions with microorganisms are pivotal for crop performance and resilience. However, it remains unclear how heritable the microbiome is with respect to the host plant genotype and to what extent host genetic mechanisms can modulate plant-microbe interactions in the face of environmental stress. Here, we surveyed the root and rhizosphere microbiome of 129 accessions of locally adapted *Zea mays*, sourced from diverse habitats and grown under control and different stress conditions. We quantified treatment and host genotype effects on the microbiome. Plant genotype and source environment were predictive of microbiome composition. Genome wide association analysis identified host genetic variants linked to both rhizosphere microbiome composition and source environment. We identified transposon insertions in a candidate gene linked to both the abundance of a keystone microbe *Massilia* and source total soil nitrogen, finding mutant plants to show a reduction in lateral root density. We conclude that locally adapted maize varieties exert patterns of genetic control on their root and rhizosphere microbiomes that follow variation in their home environments, consistent with a role in tolerance to prevailing stress.

## Introduction

Microorganisms that colonize the rhizosphere surrounding plant roots, root surfaces and internal tissues play an important role in promoting plant health and fitness under biotic and abiotic stresses (Cheng et al., 2019; Oldroyd and Leyser 2020). Specific features of the root microbiome have been shown to modify root architecture (Finkel et al., 2020), regulate nutrient homeostasis (Salas-González et al., 2020), protect against stress (Cheng et al., 2019) and impact ecosystem function (Banerjee et al., 2018). Although the overall root microbiome is largely shaped by soil properties (Bulgarelli et al., 2013), small host-mediated changes in microbiome composition can have large effects on plant fitness (Bulgarelli et al., 2012; Lundberg et al., 2012; Haney et al., 2015). Modification of crop microbiomes has been proposed as a contribution to promoting food security, while supporting a sustainable agroecosystem (de Vries et al., 2020; Singh et al., 2020). However, the extent to which host genetic mechanisms can modulate the microbiome under different environmental conditions and the genetic basis of any such control remains poorly characterized.

The diversity of traditional crop varieties (“landraces”) provides a powerful resource to investigate heritable variation in crops (Meyer and Purugganan, 2013; Cordovez et al., 2019; Raaijmakers and Kiers 2022). Furthermore, long term selection in diverse, and often challenging environments can reveal subtle signals linking plant genetic and phenotypic variation to local conditions. Maize (*Zea mays*. L) is an excellent model for investigating the genetic basis and environmental signature of plant-microbe interactions due to the extensive climatic variation across its range (Navarro et al. 2017). The domestication of maize, began 9,000 years ago when Mexican farmers started to collect the seeds of the wild grass teosinte (*Zea mays* ssp. *parviglumis*; Hake and Ross-Ibarra, 2015). During maize domestication and improvement, the root system expanded its functionality and complexity (Yu et al., 2016; Hochholdinger et al., 2018). Recent studies highlighted thatthe maize rhizosphere microbial community has also been substantially impacted by domestication (Szoboszlay et al., 2015; Brisson et al., 2019) and modern hybrid breeding (Wagner et al., 2020; Favela et al., 2021). Better understanding the genetic basis of host plant control of their microbiome and how these associations change under abiotic stress will benefit efforts promote crop resilience in the context of more sustainable agronomic practices.

Here, we profiled 3,168 root and rhizosphere microbiome samples from 129 diverse *Zea mays* accessions grown under control, nitrogen-, phosphorus- and water-limited conditions using 16S rRNA gene and ITS gene sequencing. We assessed how the native habitat of traditional varieties was predictive of root and rhizosphere microbiota assembly under our common treatments. Understanding how plant traits modulate their microbiome to enhance tolerance to environmental constraints, the extent to which this plant trait-microbe association is heritable under abiotic stresses, and how this association is encoded in the genetic program provide novel insights into establishment of beneficial host–microbiome associations. Such insights will contribute towards the generation of environment-tailored cultivars that recruit favourable microbial consortia for increasing agricultural productivity, resilience to climate change and sustainability.

## Results

### The maize microbiome responds strongly to abiotic stresses

We used 16S rRNA gene and ITS gene sequencing to characterize the root and rhizosphere microbiome of 129 *Zea* accessions, encompassing a wide range of maize and teosinte varieties. Our goal was to investigate how plant genotype, impacts crop-microbiome associations and their capacity to influence plant performance under common stress conditions. These analyses included 11 teosintes, 97 landraces, 11 maize inbred lines and 10 maize hybrids (Supplementary Fig. 1) grown in control-, low phosphorous-, low nitrogen-, and drought-treatments in a soil sourced from a long-term field experimental station (See Methods) (Supplementary Fig. 2). We sampled root and rhizosphere compartments from the first whorl of shoot-borne crown roots (Supplementary Fig. 3), in addition to collecting bulk soil. Microbial community composition differed across samples for both bacteria and fungi, with compartment (bacteria, R^2^ = 0.756; fungi: R^2^ = 0.402) explaining the largest proportion of the variation followed by stress treatment (bacteria, R^2^ = 0.052; fungi: R^2^ = 0.021) (Fig. 1a). Plant genotype (bacteria, R^2^ = 0.01; fungi: R^2^ = 0.05) was less important than either compartment or treatment (Fig. 1a). In the rhizosphere and roots, we observed significantly (One-Way ANOVA, *P* < 0.05) lower bacterial diversity under drought stress and nitrogen deficiency compared to control conditions (Supplementary Fig. 4a). In contrast, no significant differences in bacterial community diversity were observed between phosphorus deficient and control conditions (Supplementary Fig. 4a). For fungal diversity, the only significant treatment difference was (One-Way ANOVA, *P* < 0.05) lower diversity under nitrogen deficiency than control conditions in the root (Supplementary Fig. 4b). These results illustrate that both abiotic stresses and genotypes significantly explain the microbial variance although the compartment dominate the overall microbial diversity.

**Figure 1.**
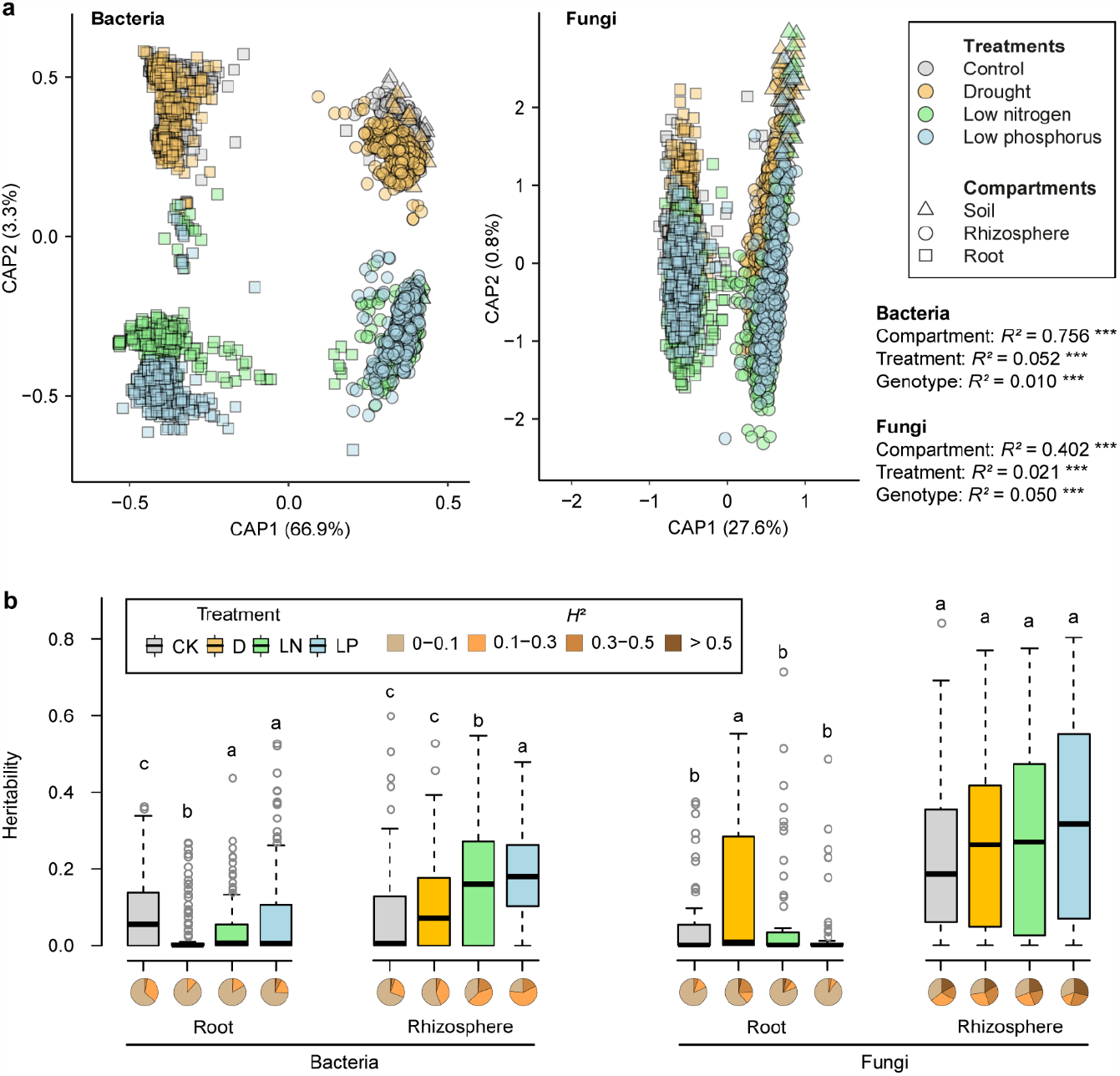
Overall diversity and heritability of microbiome among abiotic stresses. **a**, Constrained analysis of principle coordinate (CAP) ordination using Bray–Curtis dissimilarity with permutational analysis of variance (PERMANOVA) was applied to visualize significant microbiome differences across three compartments, four treatments and genotypes (*n* = 129). Datapoints for bacteria (*n* = 3138) and fungi (*n* = 3168) are color coded according to the four treatments. Compartments are shape coded. Only ASVs with reads >10 in ≥6 samples were included in the dataset. **b**, Heritability estimates of individual families under four treatments for both bacteria and fungi. The broad-sense heritability (*H*^*2*^) was calculated using highly abundant bacterial (*n* = 131) and fungal (*n* = 59) families across all samples. CK, control; D, drought; LN, low nitrogen; LP, low phosphorus. Significances are indicated among treatment groups for each compartment with Benjamini-Hochberg adjusted *P* < 0.05 (Kruskal-Wallis test, Dunn’s *post*-*hoc* test). Boxes span from the first to the third quartiles, centre lines represent the median values and whiskers show data lying within 1.5× interquartile range of the lower and upper quartiles. Data points at the ends of whiskers represent outliers. The pie charts indicate the proportional distributions of heritability frequencies.

### Keystone genera define the major differences in the microbiome

Keystone microbial taxa are defined as the drivers of microbiome structure and function (Banerjee et al., 2018). We identified putative keystone microbes among the highly abundant amplicon sequence variants (ASVs) using co-occurrence network analysis (Supplementary Datasets 1-4). Overall, the number of associations and accumulative weights of ASVs were largely positive within the bacterial or fungal networks, but negative in the inter-kingdom network (Supplementary Fig. 5; Supplementary Dataset 5). This is consistent with previous reports that inter-kingdom interactions determine the overall assembly, stability, and fitness of the root microbiome in *Arabidopsis* (Durán et al., 2018). We also observed that a high proportion of the negative inter-kingdom associations were conserved across the stress treatments (Supplementary Fig. 6; Supplementary Dataset 6). Among those, keystone taxa in the bacterial genera *Massilia, Sphingobium* and *Streptomyces* were conserved across stress treatments (Supplementary Fig. 6). Functional prediction indicates that these bacterial genera are involved in ureolysis (*Massilia*) and aerobic chemoheterotrophy (*Sphingobium* and *Streptomyces*) (Supplementary Dataset 7). The fungal keystone taxa were mainly predicted to be decomposers (37%) and pathogens (25%; Supplementary Dataset 8). Overall, our co-occurrence network analyses revealed strong negative correlations between bacterial and fungal ASVs in maize roots, while keystone bacterial members are conserved in association with other microbial members regardless of abiotic stress treatment.

### The impact of plant genotype on the rhizosphere bacterial community increases under stress

To estimate the influence of the plant genotype on microbiome composition, we estimated the correlation between the plant genetic distance matrix and the microbiome distance matrix using 97 landraces, for both root and rhizosphere. There was a significant correlation (Mantel’s statistics) between the bacterial communities and plant genotypes in both compartments (Rhizosphere, R= 0.32, *P* = 1.0e−4; Root, R= 0.23, *P* = 1.0e−4). In contrast, fungi displayed a significant correlation with the plant genotype only in the rhizosphere (R= 0.16, *P* = 0.0079) (Supplementary Fig. 7). We estimated the broad-sense heritability (*H*^2^) for the microbiome at different taxonomic levels and for individual ASVs across the experiment and then separately for each compartment and treatment combination (Supplementary Dataset 9; see methods). Across treatments, *H*^2^ was higher for the rhizosphere (*H*^2^ = 0.15; *H*^2^ = 0.14; *H*^2^ = 0.16) than the root (*H*^2^ = 0.052; *H*^2^ = 0.049; *H*^2^ = 0.052) at the level of families (Fig. 1b), genera (Supplementary Fig. 8a) or ASVs (Supplementary Fig. 8b), respectively. Nutrient stress significantly (Kruskal-Wallis test, Dunn’s *post*-*hoc* test with BH adjusted *P* < 0.05) increased *H*^2^ (control, *H*^2^ = 0.078; low nitrogen, *H*^2^ = 0.16; low phosphorus, *H*^2^ = 0.18) for the bacterial rhizosphere microbiome, but not of the fungal microbiome. To identify plant genetic loci affecting microbiome composition, we performed genome-wide association (GWA) analysis for the relatively high heritable (*H*^2^ > 0.1) microbial traits at the level of overall diversity, families, genera and ASVs (Supplementary Dataset 10). We did not recover significant markers in association with overall measures of microbial alpha-diversity (Shannon index) (Supplementary Dataset 11). We did, however, identify significant associations with individual ASVs (Supplementary Dataset 10). Overall in our experiment, these data indicate an increasing impact of the plant genotype on microbiome composition, especially the composition of the rhizosphere bacterial community under stress.

### Plant source habitats predict the root and rhizosphere microbiome

To address the hypothesis that variation in plant modulation of the root microbiome is a reflection of differences in native environment, we assessed the potential of environmental descriptors for the point of collection to predict the microbiome in our standardized growth chamber experiments (Supplementary Fig. 1; Supplementary Dataset 12). To reduce the complexity of the microbiome data, we used Spearman correlation analysis to define four microbial assemblies corresponding to dominant ASVs (Supplementary Figure 9). We then sought evidence of covariation among microbial assemblies and environmental descriptors (Supplementary Figure 10). We used structural equation modeling to quantify the cumulative effects of source environment, plant genetic diversity, stress treatment, domestication status and biomass on the four microbial assemblies. These analyses demonstrated an impact of plant genotype and source environment on specific assemblies of microbiome. Low nitrogen treatment, source mean annual temperature, source precipitation and plant genotype strongly impacted the microbiome assemblage (Supplementary Figure 11), one notable example being the abundance of the genus *Massilia*, which belongs to the previously mentioned *Oxalobacteraceae* (Supplementary Figure 12). Overall, prediction was better for bacterial data than for fungal data, and better for rhizosphere than root (Fig. 2a; Supplementary Fig. 13). Interestingly, microbiome composition could be predicted more accurately with environmental descriptors or a combination of these with plant genetic markers than with genetic markers alone (Fig. 2a; Supplementary Fig. 14−16). Under the conditions of our experiment, ecological modelling and prediction analyses show potential effects of source environment of locally adapted maize on the composition of the rhizosphere communities.

**Figure 2.**
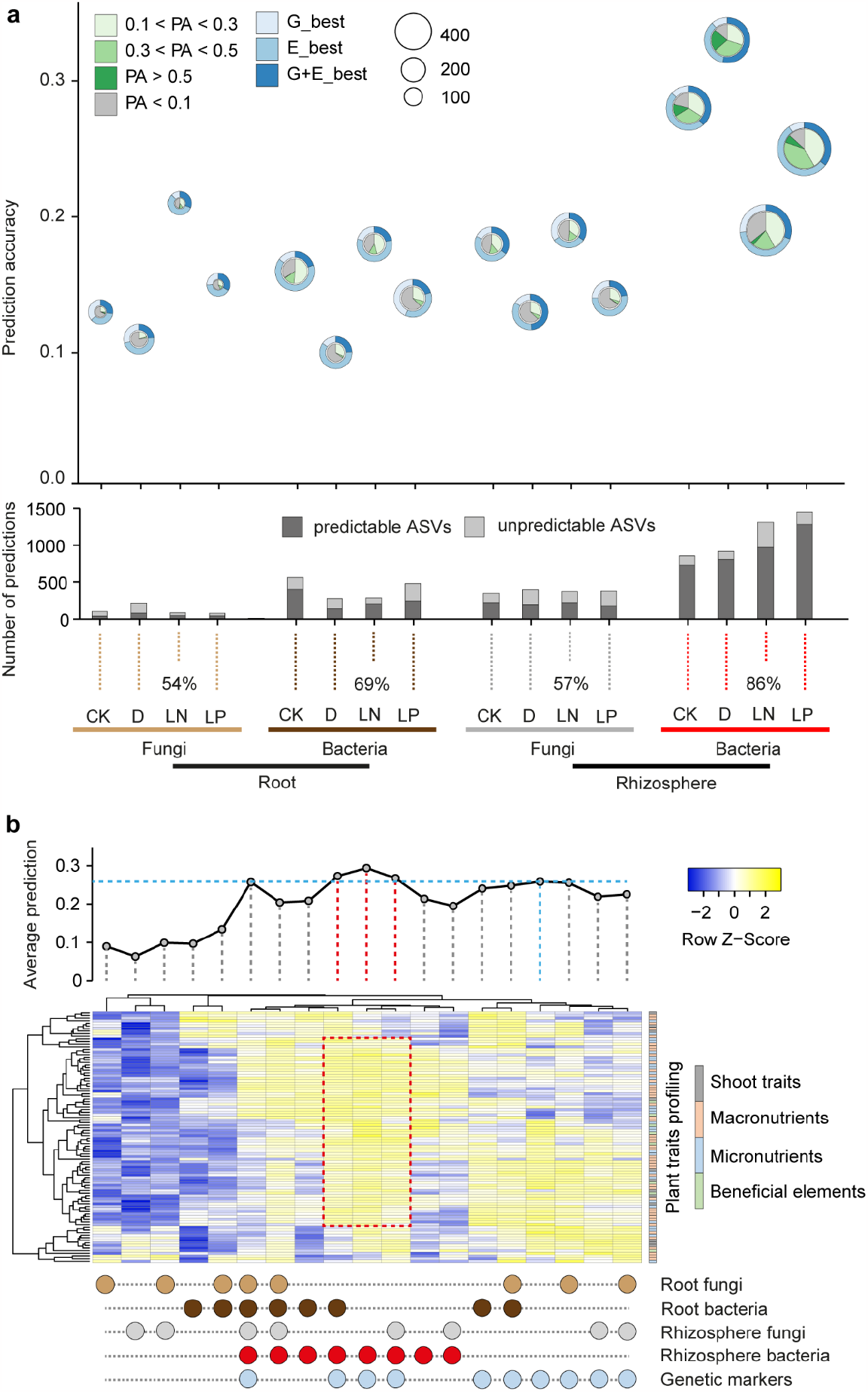
Genomic, environmental and microbial prediction of host-microbe interactions and plant traits. **a**, Microbiome traits prediction using genetic markers and environmental characters. Inner pie charts describe the proportion of ASVs with four different magnitudes of prediction accuracies from different treatments across compartments. Outer circles define the best prediction patterns observed by applying the genetic markers (G_best) alone, environmental characters (E_best) alone or combined genetic markers and environmental characters (G+E_best). The numbers denote the average prediction accuracies for microbial ASVs from different treatments across compartments. Only ASVs with heritability (*H*^*2*^) >0.1 were considered in prediction analysis. PA, prediction accuracy. Bar plots indicate the proportions of predictable (PA >0.1) and unpredictable (PA <0.1) ASVs from the total predictions. CK, control; D, drought; LN, low nitrogen; LP, low phosphorus. **b**, Plant traits prediction using genetic markers and microbiome traits. A curved line describes the average prediction accuracy for plant traits using microbiome data alone, genomic data alone or combined genomic and microbiome traits data. A heatmap illustrates the standardized prediction accuracy for fitness traits across different microbiome features combined with genetic markers. Shoot traits include the biomass, leaf area and chlorophyll measured by SPAD value. Nutrient uptake properties include the concentration and content of macronutrients (nitrogen, phosphorus, potassium, calcium, magnesium and sulfur), micronutrients (iron, manganese, zinc and boron) and beneficial elements (aluminium and sodium).

### Consideration of the rhizosphere bacterial community improves prediction of plant traits

To assess the relationship between the microbiome and plant growth and physiology, we used a two-step strategy combining genomic prediction and Random Forest models based on environmental descriptors. First, we compared prediction of plant growth and nutrient accumulation traits using plant genotype alone or in combination with microbiome ASVs abundance. The combination of plant genotype and rhizosphere bacterial community composition provided the highest average prediction ability (29%) (Fig. 2b; Supplementary Datasets 13 and 14). We confirmed this result employing an alternative approach to fit a ridge regression mixed model, observing ∼10%−15% increase of prediction accuracy when using both genetic and microbiome data (Supplementary Figure 17). As has been previously seen in foxtail millet (Wang et al., 2022), we showed a conserved pattern that the rhizosphere microbiome combined with genotype data increased the average prediction accuracy ∼7% of 12 agronomic traits compared to genetic markers alone (Supplementary Figure 18). We then explored relationships among source environments, genetic differentiation and specific microbial taxa. As a measure for the pattern of similarity among samples, we calculated matrices of pairwise distance using the observed microbiome ASVs in different treatments, and two source environmental descriptors (*elevation* and *geographical distance*). Mantel tests were used to study the correlations between different distance matrices. We observed that the correlation between the rhizosphere microbiome and source environment was higher than that between the root microbiome and environment. On average, the correlations of inter-treatment and treatment-environment similarity patterns as characterized by bacterial communities were higher than by fungal communities (Supplementary Fig. 19). To reduce dimensionality, we extracted the first five principal components (PCs) from the microbiome ASV data. We then used a Random Forest (RF) approach to predict these PCs using different environmental descriptors as explanatory variables (Supplementary Dataset 12). We observed the highest accuracy for the rhizosphere bacteria PC2 (Supplementary Fig. 20a) using environmental predictors including *photosynthetically active radiation* and *potential evapotranspiration* (Supplementary Fig. 20b). Prediction of individual ASVs was less successful (Supplementary Fig. 21), although significant predictors were identified for specific examples belonging to the *Oxalobacteraceae*, including *Massilia* (Supplementary Fig. 22). These results suggest that source environment plays effect on plant genetic variation in regulation of the microbiome composition with an impact of plant traits.

### A candidate gene linked to source environment associates with *Oxalobacteraceae* abundance and root branching

Across our samples, we detected five highly abundant bacterial families (*Pseudonocardiaceae, Streptomycetaceae, Chitinophagaceae, Oxalobacteraceae* and *Comamonadaceae*; Fig. 3a), and three highly abundant fungal families (*Aspergillaceae, Trichocomaceae* and *Nectriaceae*; Supplementary Fig. 23). In particular, the bacterial taxon *Oxalobacteraceae* is the only family under nitrogen limitation showed the highest *H*^2^ among all families in our experiment (Fig. 3b; Supplementary Dataset 15). *Oxalobacteraceae* have been previously proposed to play an important role in maize tolerance to nitrogen limitation when grown in nitrogen-deficient soils (Yu et al., 2021). To identify loci associated with variation in the microbiome and differences in source environment, we used our RF models to predict *Oxalobacteraceae* ASVs for 1781 previously genotyped traditional varieties (Navarro et al. 2017) on the basis of associated source environmental descriptors and subsequently implemented GWA analyses (Fig. 4a). One of the best predictions (RF model R^2^ = 0.28) was for root abundance of ASV37, belonging to the genus *Massilia* (*Oxalobacteraceae*), in the low nitrogen treatment, consistent with our previous estimates of *H*^2^. Collectively, GWA hits from environmental predictions of ASV37 abundance for the 1781 panel overlapped more than expected by chance with the hits from the observed ASV37 data in the smaller 129 panel (Supplementary Fig. 24). The top GWA hit for predicted ASV37 root abundance under low nitrogen (SNP S4_10445603) fell within the gene Zm00001d048945 on chromosome 4 (Fig. 4a and b; Supplementary Dataset 16). Across the 1781 panel, the minor allele at SNP S4_10445603 was more abundant at higher predicted ASV37 abundance but lower source soil nitrogen content (Fig. 4c). These findings are consistent with a specific gene contributing to the geographical adaptation to nitrogen-poor soil by facilitating enhanced association with *Massilia* (Yu et al., 2021; Supplementary Fig. 25). The gene Zm00001d048945 is most strongly expressed in the root cortex (Fig. 4d; https://www.maizegdb.org/gene_center/gene/Zm00001d048945) and is predicted to encode a TPX2 domain containing protein related to the WAVE-DAMPENED2 microtubule binding protein that functions in *Arabidopsis* root development (Yuen et al., 2003) and lateral root initiation (Qian et al., 2022). Using root architectural data available for the 97 landraces, we found a positive correlation between lateral root density and ASV37 abundance (*r* = 0.2, *P* = 0.03; Fig. 4e). To test the hypothesis that variation in Zm00001d048945 contributes to a root-architecture-related effect on ASV37, we identified transposon insertional mutants in two different genetic backgrounds (B73 and F7; Supplementary Fig. 26). Plants homozygous for transposon insertions in Zm00001d048945 showed a significant reduction in lateral root density (Fig. 4f and g). We interpret these results as evidence that variation at Zm00001d048945 alleles, affect root traits and that this variation also affects *Massilia* abundance in the root under nitrogen limitation.

**Figure 3.**
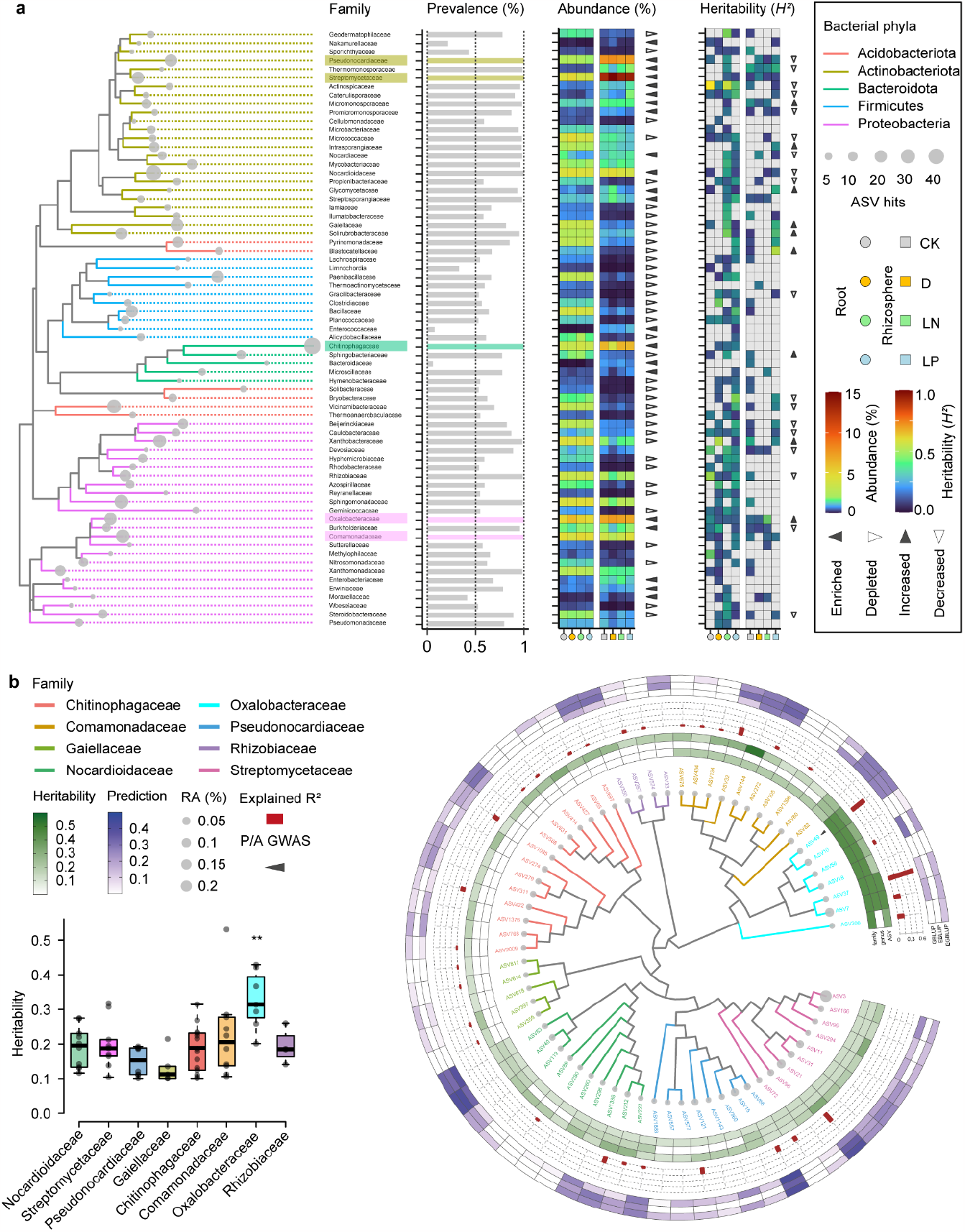
Dominated and heritable bacterial families of maize root and rhizosphere microbiome under abiotic stresses. **a**, Maximum-likelihood phylogeny of dominant bacterial families (n > 5). Circle sizes along the branches of the tree indicate the number of ASVs observed in association with microbial families. Colour coded families are clustered at the phylum level. Bar plots describe the prevalence according to the proportional sample size. The heatmaps illustrate the standardized mean relative abundance and the estimated heritability of microbial families from the root to the rhizosphere. Triangles represent the enrichment or depletion of microbial families, and increased or decreased heritability from the root to the rhizosphere. The significance levels were controlled at two levels (*: *p* <0.05; **: *p* <0.01). **b**, Phylogenetic tree of dominant bacterial ASVs (*n* = 126) of roots grown under nitrogen-poor condition. Dot size corresponds to relative abundance. Inner heatmap from inside to outside indicates heritability (*H*^*2*^ >0.1) at the family, genus and ASV level. Red bar plots describe the explained variance by GWAS. The outer heatmap indicates the predictions by genomic best linear unbiased prediction (GBLUP), or based on the environmental best linear unbiased prediction (EBLUP) or prediction based on both genomics and environment (EGBLUP). Triangles indicate significant associations with the presence/absence (P/A) GWAS. Color coded tree branches of ASVs are clustered at the family level. Box plot indicates significantly higher heritability of *Oxalobacteraceae* compared to other families.

**Figure 4.**
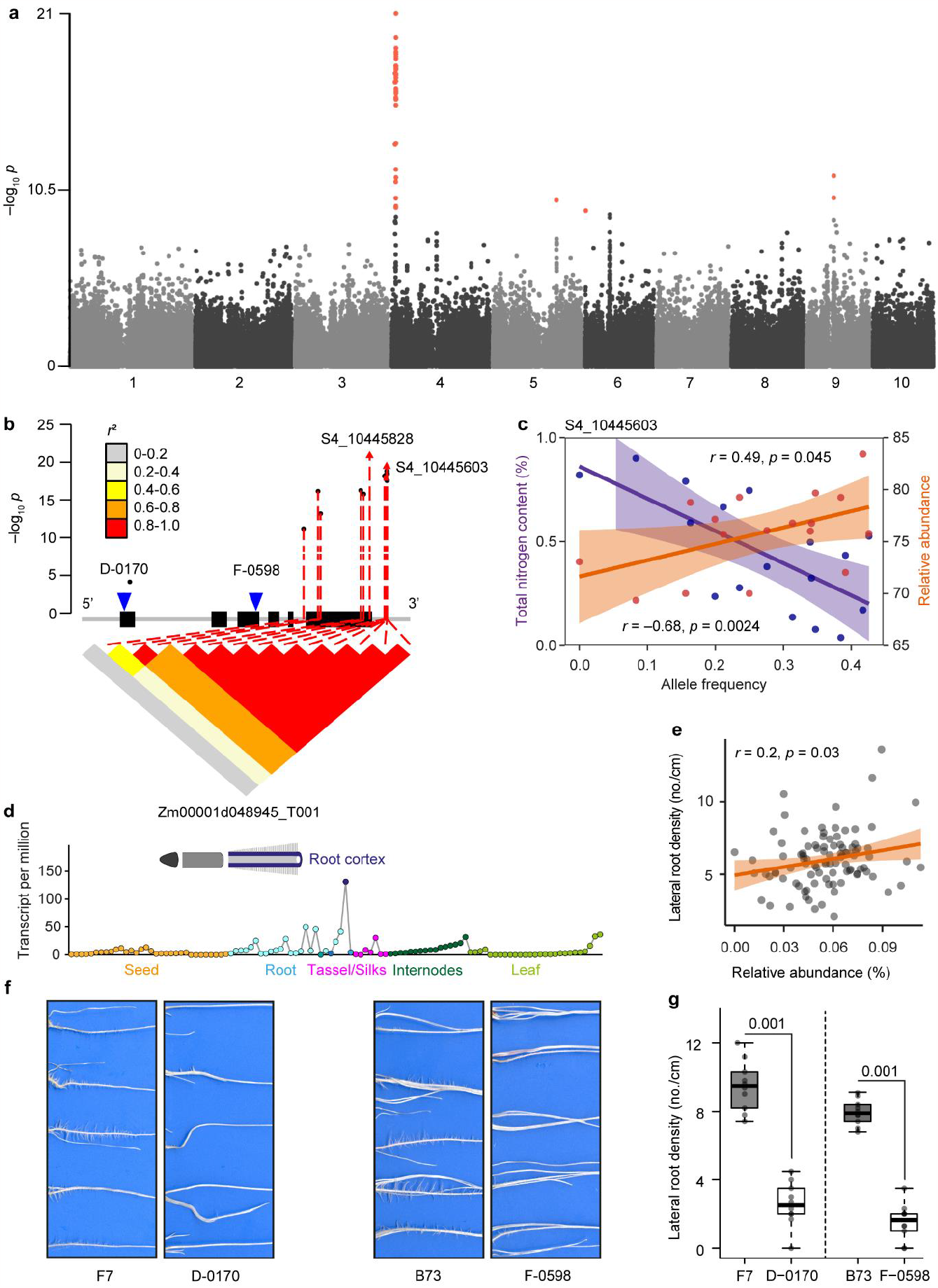
Environmental selection facilitates microbiome-driven root phenotypic association with to nitrogen availability of source habitats. **a**, Manhattan plots showing environmental GWAS of adaptive of specific *Massilia* ASV37. **b**, Linkage disequilibrium (LD) plot for SNPs within 2.5kb of gene Zm00001d048945. Exons in the gene model are indicated by black bins. All significant SNPs are linked (red) to the LD plot (*P* < 1.0 × 10^−7^). Arrows indicate the positions of the peak SNPs. The colour key (grey to red) represents linkage disequilibrium values (*r*^2^). Blue triangles indicate the transposon insertion positions of the two mutant alleles D-0170 and F-0598. **c**, Pearson correlation coefficient analysis of allele frequency (S4_10445603) with soil total nitrogen content (purple) and predicted relative abundance of ASV37_Root_LN (orange) across 1781 geographical locations worldwide. **d**, Tissue-specific expression of gene Zm00001d048945 according to the eFP Browser database. **e**, Pearson correlation coefficient analysis of lateral root density with relative abundance of ASV37_Root_LN (orange) among 97 maize landraces. Scatter plots show best fit (solid line) and 95% confidence interval (colour shading) for linear regression. **f** and **g**, Root phenotypes and lateral root density of two independent Mu-transposon insertion mutant alleles in comparison to the corresponding wild types (B73 and F7). Significances are indicated between wild type and mutant for different genetic backgrounds (two-tailed Student’s *t*-tests). Boxes span from the first to the third quartiles, centre lines represent the median values and whiskers show data lying within 1.5× interquartile range of the lower and upper quartiles. Data points at the ends of whiskers represent outliers.

## Discussion

During domestication plants have developed high productivity and environmental resilience, but may have also lost beneficial microbiome-associated traits compared with their wild relatives (Haney et al., 2015; de Vries et al., 2020). Thus, bringing back important plant traits supporting beneficial microbes from wild relatives and broader crop diversity may contribute to adaptation of crops to future climatic challenges. In this study, we investigated the host-microbiome association and tried to understand whether and how source environments of traditional varieties relate to microbiome assembly under multiple abiotic stresses in maize. Examination of microbiomes across diverse germplasm demonstrated that plant genotype significantly impacts the microbiome, more so under abiotic stresses. Our genetic and environmental analyses support the hypothesis that plant genetic variation impacts microbiome assembly in crops (Deng et al., 2020; Escudero-Martinez et al., 2022; Meier et al., 2022; Oyserman et al., 2022; Wang et al. 2022). Rhizosphere microbial diversity supports rhizosphere function under harsh environments (Ramirez et al., 2018) and is heritable trait across environments (Walters et al., 2018). We report here a significant improvement in plant trait prediction when combining rhizosphere microbiome with plant genetic data. Binominal regression and correlation analyses between microbial traits and source environmental variables among traditional varieties suggest that microbiome assemblage may contribute to beneficial plant trait-microbe association underlying stress-resilience.

Although environmental conditions were dominant drivers of the crop microbiome, we found certain microbial taxa that were consistently influenced by genetic variability in maize, and whose abundance correlated with plant traits. The endogenous genetic program that underlies root development can coordinate microbiome assembly and plant mineral nutrient homeostasis (Salas-González et al., 2020). Notably, we found that environment-associated alleles may promote root differentiation and microbiome-driven nitrogen deficiency tolerance. These results provide strong support for a genetic basis for variation in the abundance of the bacterial taxon *Massilia* (*Oxalobacteraceae*) under nitrogen deficiency, illustrating the importance of specific bacteria for root development (Finkel et al., 2020), nitrogen nutrition (Zhang et al., 2019) and reciprocal interaction (Yu et al., 2021). Taken together, this study advances the current understanding of the plant-trait-microbiome interactions that connecting genetic variation to microbiome composition among a broad array of maize and their relatives in multiple environmental treatments, as well as identifying a specific gene with a compelling association with both the environment and a bacterial taxon *Massilia*. These findings help to close the knowledge gap between how plants impact the soil microbiome and how this functional interaction of the microbiome can be translated into crop resilience to nutrient limitation.

## Material and Methods

### Plant material, soil collection and growth conditions

The germplasm used in this study was selected to represent a broad diversity ranging from the maize progenitor teosinte to local open pollinating landraces and modern inbred lines and hybrids (Supplementary Dataset 17; Supplementary Fig. 1). We obtained the 11 geographically diverse teosinte accessions from the North Central Regional Plant Introduction Station (NCRPIS) and the International Maize and Wheat Improvement Center (CIMMYT). Moreover, we received the 97 landrace accessions from NCRPIS and these accessions were derived from the ten American countries which cover the major domestication areas of maize (Supplementary Fig. 1a). The modern breeding germplasm includes seven genetically diverse inbred lines (Baldauf et al., 2018) covering the major heterotic groups stiff-stalk and non-stiff stalk and four additional tropical inbred lines (Supplementary Fig. 1b). We have produced the ten hybrids by crossing the ten inbred lines with the reference inbred line B73 as the common mother plant (Supplementary Fig. 1c). Soil used for phytochamber pot experiments was dug from the Dikopshof long-term fertilizer field experiment established in 1904 near Cologne, Germany (50°48′21′′N, 6°59′9′′E) (Supplementary Fig. 2a). In this study, we collected soil subjected to three different fertilization managements including control soil fertilized with all nutrients, low nitrogen soil fertilized without nitrogen and low phosphorus soil fertilized without phosphorus as defined by Rueda-Ayala et al. 2018. The general soil type is classified as a Haplic Luvisol derived from loess above sand. Approximately the first 0-20 cm of the soil were collected and placed in a clean plastic bag. Subsequently, collected soil was dried at room temperature in clean plastic trays for about one week and sieved with a 4 mm analytical sieve (Retsch, Haan, Germany) to remove stones and vegetative debris. The sieved soil for the whole experiment was then homogenized with a MIX125 concrete mixer (Scheppach, Ichenhausen, Germany) (Supplementary Fig. 2a). The air-dried soil was ground into powder for the analysis of carbon, nitrogen, phosphorus and five metal elements (K, Fe, Mn, Cu, Zn). Soil pH was measured in deionized water (soil: solution ratio, 1:2.5 w/v) using a pH-meter 766 (Knick, Berlin, Germany). The basic physical and chemical properties of these soils are provided in Supplementary Table 1.

Local landraces are open-pollinated varieties and can vary largely on seed traits. Therefore, we covered a broad geographic area but also confirmed the homogeneity of the 97 landraces concerning seed size, seed color, and seed quality prior our phytochamber experiments (Supplementary Fig. 2b). Seeds were surface-sterilized with 6% NaClO for 10 min, and rinsed 3 times with sterile deionized water to eliminate any seed-borne microbes on the seed surface. The sterilized seeds were pre-germinated for 3 days in a paper roll system using germination paper (Anchor Paper Co., St. Paul, MN, USA) with sterile deionized water. Then seedlings with primary roots of ca. 1–2 cm length were transferred to soil-filled pots (7 × 7 × 20 cm^3^) in a 16/8-h light/dark, 26/18 °C cycle and were grown for 4 weeks in a walk-in phytochamber. A detailed sowing and transfer plan is provided in Supplementary Fig. 2c. No additional fertilizer was added.

### Experimental design and treatments

The experiment was performed in a split plot design with three replications comprising four stress treatments on the main plots (trays) (Supplementary Fig. 27), e.g. fully fertilized control (CK) soil, no nitrogen fertilized low nitrogen (LN) soil, no phosphorus fertilized low phosphate (LP) and CK soil with drought (D) treatment. As controls, we used six pots without plants as ‘bulk soil’ samples (B), which were distributed across the main plots. Each tray contained a similar number of pots (subplots) with the different genotypes and bulk soil. The three replicates were performed at three different periods in the same growth chamber (Supplementary Fig. 27). For each stress treatment, we generated an alpha design for the genotypes and controls with three replicates and four incomplete blocks per replicate. The incomplete blocks were assigned to trays and replicates corresponded to the three replications of the experiment in time. To facilitate watering, pots subjected to the same treatment were allocated on the same tray. These trays were further randomized in the chamber. Distribution of all pots in each tray were randomized using a true random generator (excel function “RAND”), and trays were reshuffled every week in the growth chamber without paying attention to the pot labels. Since soil water availability will significantly affect the harvest of the rhizosphere and initiation of crown roots, we have performed a preliminary experiment with different water regimes (i.e. 33%, 22%, 17% water holding capacity) to ensure the establishment of suitable drought conditions and to facilitate rhizosphere harvesting and the optimal formation of the different whorls of crown roots (Supplementary Fig. 2c and 28). In brief, different volumes of sterilized water e.g. 60 ml, 40 ml, 30 ml were mixed with 500 g dry soil by spraying water and were then homogenized with a 4 mm sieve (Retsch). Each water regime was maintained by spraying water to the soil surface according to the weight loss of water during the 4-week culture. Plant height, total leaf area, shoot and root fresh biomass from the representative genotypes B73 and Mo17 were recorded. Moreover, the multifunctional device COMBI 5000 (STEP Systems, Nuremberg, Germany) was used to measure soil variables e.g. soil moisture and electronic conductivity directly in each soil pot during each experimental run. The covariates including sample harvest time, ID of person performing DNA extraction together with the determined soil variables were collected and used for downstream data analysis (Supplementary Dataset 18).

### Characterization of native collection sites of maize landraces

Geographical coordinates and elevation information of the collection sites for maize landraces were retrieved from the public database of the U.S. National Plant Germplasm System (https://www.grin-global.org/) and provided in Supplementary Dataset 17. Most of the landraces were received in the years 1980-1994 and were maintained by NCRPIS. To get the climate and soil variables based on the geographical coordinates for each site, we first compiled climatic and soil descriptors representative of the long-term averages of their point of origin, following methods in Lasky et al. 2015. All used databases are publicly available and have global coverage. Data was collected from WorldClim (Zomer et al. 2008), the NCEP/NCAR reanalysis project (https://psl.noaa.gov/data/reanalysis/reanalysis.shtml) (Kalnay et al., 1996), NASA SRB (https://asdc.larc.nasa.gov/project/SRB), Climate Research Unit (CRU) (New et al. 2002), SoilGrids (Hengl et al. 2017) and the Global Soil Dataset (GSD) (Shangguan et al. 2014). All 156 bioclimatic and soil variables were merged with the maize germplasm identity into the Supplementary Dataset 12. The related information of total soil nitrogen, available phosphorus, and annual precipitation are provided in the Supplementary Fig. 29.

### Determination of shoot phenotypic traits and ionome profile

Aboveground phenotypic traits were determined for all 129 genotypes on the day of harvest in the phytochamber. The leaf area and chlorophyll index as measured by SPAD were determined as described accordingly (Yu et al., 2021) and are provided in Supplementary Dataset 19. The complete aboveground part of maize plants excluding the seed was harvested and heat treated at 105 °C for 30 min, dried at 70 °C to constant weight, weighed as the shoot dry biomass and then ground into powder. Approximately 6 mg of ground material was used to determine total nitrogen concentration in an elemental analyzer (Euro-EA, HEKAtech). Data were then calculated into peak areas by the software Callidus, providing quantitative results using reference material as a calibration standard. The same plant material was used to determine the concentrations of 13 additional mineral nutrients. In brief, approximately 200 mg of same ground material was weighed into polytetrafluoroethylene digestion tubes, and concentrated nitric acid (5 ml, 67–69%; Bernd Kraft) was added to each tube. After 4 h of incubation, samples were digested under pressure using a high-performance microwave reactor (Ultraclave 4, MLS). Digested samples were transferred to Greiner centrifuge tubes and diluted with deionized (Milli-Q) water to a final volume of 8 ml. Element analysis was carried out by Inductively Coupled Plasma-Optical Emission Spectroscopy (iCAP 7400 duo; Thermo Fisher Scientific). For sample introduction a SC-4 DX autosampler with prepFAST Auto-Dilution System (ESI, Elemental Scientific) was used. A three-point external calibration curve was set from a certified multiple-standards solution (Custom Multi-Element Standard_PlasmaCAL, S-prep GmbH). The element Yttrium (ICP Standard Certipur®, Merck) was infused online and used as internal standard for matrix correction. All ionome data including concentrations and contents of all mineral nutrients are provided in the Supplementary Dataset 20.

### Root and rhizosphere samples harvest for microbiome analysis

The root and rhizosphere samples collection were performed from 4-week-old maize plants as previously described (Yu et al., 2021). In brief, whole root systems were carefully taken out from each pot and vigorously shaken to remove all soil not firmly attached to the roots. During this stage, most genotypes have consistently started to form the 2^nd^ whorl of shoot-borne crown roots with a length of 3-10 cm. To synchronize the harvest for precise comparisons among genotypes, we collected the fully developed 1^st^ whorl of shoot-borne crown roots initiated from the coleoptilar node for all maize genotypes (Supplementary Fig. 3a). These crown roots were identified similarly developmental status with mature lateral roots. Two dissected crown roots with tightly attached soil were placed into a 15 ml Falcon (Sarstedt) tube and immediately frozen in liquid nitrogen and stored at -80 °C before extraction of rhizosphere soil. The rhizosphere samples were defined and extracted into PowerBead tubes (Mo Bio Laboratories) as described previously (Yu et al., 2021). The root samples were harvested from another crown root from the same plant that immediately washed by tap water and rinsed with three times of sterilized water followed by tissue drying and placed in PowerBead tubes (Supplementary Fig. 3b). Sample processing steps for root and rhizosphere have been performed by a designated person to avoid systematic errors. The bulk soil samples were also collected from the unplanted pots. DNA extractions were performed soon after root and rhizosphere samples were harvested, following the PowerSoil DNA isolation kit (Mo Bio Laboratories) protocol.

### Amplicon library preparation and sequencing

Amplicon library construction was processed with a similar workflow as previously described (Yu et al., 2021) (Supplementary Fig. 3c). In brief, for bacterial 16S rRNA gene libraries, the V4 region was amplified using the universal primers F515 (5′ GTGCCAGCMGCCGCGGTAA 3′) and R806 (5′ GGACTACHVGGGTWTCTAAT 3′) (Caporaso et al. 2011). For fungal amplicon sequencing, the *ITS1* gene was amplified by the primer pair F (5′ CTTGGTCATTTAGAGGAAGTAA 3′) and R (5′ GCTGCGTTCTTCATCGATGC 3′). PCR reactions were performed with Phusion High-Fidelity PCR Master Mix (New England Biolabs) according to the manufacturer’s instructions. Subsequently, only PCR products with the brightest bands at 400-450 base pairs (bp) were chosen for library preparation. Equal density ratios of the PCR products were mixed and purified with the Qiagen Gel Extraction Kit. Sequencing libraries were generated using the NEBNext Ultra DNA Library Pre Kit for Illumina, following the manufacturer’s recommendations and with the addition of sequence indices. The library quality was checked on a Qubit 2.0 Fluorometer (Thermo Scientific) and Agilent Bioanalyzer 2100 system. Finally, the qualified libraries were sequenced by 250-bp paired-end reads on a MiSeq platform (Illumina).

### 16S rRNA gene and ITS gene sequence processing

Raw sequencing reads were processed following a similar workflow as previously described (Yu et al. 2021). Briefly, paired-end 16S rRNA amplicon sequencing reads were assigned to samples based on their unique barcode and truncated by cutting off the barcode and primer sequence. Paired-end reads were merged using FLASH (v1.2.7) (Magoč and Salzberg 2011) and the splicing sequences were called raw tags. Sequence analyses were performed by QIIME 2 software (v2020.6) (Bolyen et al. 2019). Raw sequence data were demultiplexed and quality filtered using the q2-demux plugin followed by denoising with DADA2 (Callahan et al. 2016) (via q2-dada2). Sequences were truncated at position 250 and each unique sequence was assigned to a different ASV. Taxonomy was assigned to ASVs using the q2-feature-classifier (Bokulich et al. 2018) and the classify-sklearn naïve Bayes taxonomy classifier against the SSUrRNA SILVA 99% OTUs reference sequences (v138) (Yilmaz et al. 2014) at each taxonomic rank (kingdom, phylum, class, order, family, genus, species). Mitochondria- and chloroplast-assigned ASVs were eliminated. Out of the remaining sequences (only features with >10 reads in ≥2 samples) were kept to build an ASV table. In order to study phylogenetic relationships of different ASVs, multiple sequence alignments were conducted using mafft (via q2-alignment) (Katoh et al., 2002) and the phylogenetic tree was built using fasttree2 (via q2-phylogeny) (Price et al., 2010) in QIIME 2. Those sequences that did not align were removed. ITS amplicon data were processed the same as 16S amplicon data except that used the UNITE 99% ASVs reference sequences (v10.05.2021) (Abarenkov et al., 2021) to annotate the taxonomy.

### Statistical analyses for microbial community assembly

In consideration of experimental design, here we treated the trays as the main plots for different treatments as a random effect. There were four trays per period/replicate, and a replicate effect was considered to account for differences between the three replicates. All downstream analyses were performed in R (v4.1.0) (R Core Team, 2021). Briefly, ASV tables were filtered with ≥10 reads in ≥2samples. For α-diversity indices, Shannon index was calculated using ASV tables rarefied to 1,000 reads. For all the following analyses ASVs which express ≤0.05% relative abundance within ≤5% samples were filtered. After filtering taxa, the samples with ≤1000 reads were also removed. Bray– Curtis distances between samples were calculated using ASV tables that were normalized using ‘varianceStabilizingTransformation’ function from DESeq2 (v1.34.0) package (Love et al., 2014) in R. Constrained ordination analyses were performed using the ‘capscale’ function in R package vegan (v2.5-7) (Oksanen et al., 2020). To test the effects of compartment, treatment and genotype on the microbial composition community, variance partitioning was performed using Bray–Curtis distance matrix between pairs of samples with a permutation-based PERMANOVA test using ‘adonis’ function in R package vegan (Oksanen et al., 2020).

### Inter-kingdom associations by network analysis

The method SPIEC-EASI (SParse InversE Covariance Estimation for Ecological Association Inference) implemented in SpiecEasi (v1.1.2) R package was used to construct the inter-kingdom microbial associations (Kurtz et al., 2015) and network was visualized by Cytoscape (v3.9.1). For this network inference, only ASVs with relative abundance >0.05% in ≥10% samples were used. The filtered bacterial and fungal ASV table were combined as the input followed by the default centered log-ratio (CLR) transformation. The neighborhood selection measured by the Meinshausen and Bühlmann (MB) method (Meinshausen and Bühlmann 2006) was selected as the inference approach. The number of subsamples for the Stability Approach to Regularization Selection (StARS) was set to 99.

### Genotyping of 129 maize genotypes

Genomic DNA was extracted from leaves of bulked maize seedlings subjected to different treatments and replicates for each genotype (Supplementary Fig. 3). The genetic variation across the maize genotypes was characterized using a GenoBaits Maize40K chip containing 40 K SNP markers, which was developed using a genotyping by target sequencing (GBTS) platform in maize (Guo et al., 2019). In brief, DNA fragmentation, end-repair and adding A-tail, adapter ligation and probe hybridization were performed. After ligation of the adapters and clean up, fragment size selection was done with Beckman AMPureBeads and a PCR step to enrich the library. Quantity and quality of the libraries were determined via Qubit™ 4 Fluorometer (Invitrogen) and Agilent 2100 Bioanalyzer, respectively. In total, 129 qualified and enriched libraries were sequenced as 250-400 bp on an MGISEQ-2000 (MGI, Shenzhen, China). The quality of raw sequencing reads was assessed and filtered by fastp (version0.20.0, www.bioinformatics.babraham.ac.uk/projects/fastqc/) with the parameters (-n 10 -q 20 -u 40). The clean reads were then aligned to the maize B73 reference genome v4 using the Burrows-Wheeler Aligner (BWA) (v0.7.13, bio-bwa.sourceforge.net) with the MEM alignment algorithm. The SNPs were then called using the UnifiedGenotyper tool from Genome Analysis Toolkit (GATK, v3.5-0-g36282e4, software.broadinstitute.org/gatk) SNP caller. The genetic distance matrix was calculated based on pairwise Rogers’ distance (Rogers 1972). A principal component analysis (PCA) was performed based on the filtered SNPs by GCTA software (Yang et al., 2011). A phylogenetic tree (Supplementary Fig. 30) was generated using the neighbour-joining method as implemented in Mega 10.0.4 with 1,000 bootstraps using MEGA-X (Kumar et al., 2018).

### Analyses of phenotypic data

For the three fitness phenotypes (SPAD, leaf area and biomass), we first performed the outlier test using the following model for a given stress treatment:

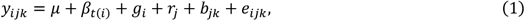

where *y*_*i jk*_ is the observation of the *i*-th genotype in the *k*-th block of the *j*-th complete replicate. *μ* is the general mean, *β*_*t(i)*_ is the effect of the *t*(*i*)-th subpopulation (*t*(*i*) indicates the subpopulation that the *i*-th genotype belongs to. There are four subpopulations: teosinte, landraces, inbred lines and hybrids.), *g* _*i*_ is the effect of the *i*-th genotype, *r*_*j*_ is the effect of the *j*-th replicate, *b*_*jk*_ is the effect of the *k*-th block nested within the *j*-th replicate and *e*_*i jk*_ is the residual term. All effects except the general mean were assumed to be random and follow an independent normal distribution.

After fitting the model, the residuals were standardized by the rescaled median of absolute deviation from the median (MAD) and then a Bonferroni-Holm test was applied to flag the outliers (Bernal-Vasquez et al., 2016).

For all traits including fitness phenotypes and microbial traits, we estimated the broad-sense heritability (also referred as repeatability in this case) in each treatment. The following model was used to estimate the heritability:

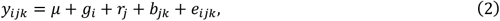

where all notations were the same as in (1).

The heritability was calculated using the following formula:

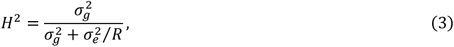

where 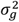 and 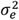 are the estimated genotypic and residual variance, *R* is the number of replications.

The best linear unbiased estimations (BLUEs) of all genotypes for each trait in each treatment were obtained by fitting Model (2) once more, assuming the general mean and genotypic effects are fixed and all other effects are random. All linear mixed models were fitted using the software ASReml-R 4.0 (Butler et al., 2017).

### Statistical framework for GWAS

Prior to GWAS, we first performed quality control for the genotypic data. In brief, the missing genotypic values were imputed using the software Beagle 5.2 (Browning et al., 2018). After imputation, we removed the markers with minor allele frequency (MAF) <0.05. As heterozygous loci were very common in our data set, we also removed markers whose maximum genotype frequency is >0.95. In total, 157,785 SNP markers were used for GWAS. For all traits, GWAS was performed separately for each treatment (i.e., using the BLUEs within the treatment as the response variable). For microbiome ASVs and alpha-diversity traits, only those with a heritability >0.1 were used for GWAS.

A standard “Q+K” linear mixed model (Yu et al., 2006) was used in GWAS. More precisely, the model is of the following form:

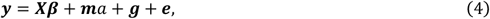

where ***y*** is the *n*-dimensional vector of phenotypic records (i.e. BLUEs within a certain treatment, *n* is the number of genotypes), ***β*** is the *k*-dimensional vector of fixed covariates including the common intercept and the subpopulation effects. ***X*** is the corresponding *n* × *k* design matrix allocating each genotype to the subpopulation it belongs to. *a* is the additive effect of the marker being tested, *m* is the *n*-dimensional vector of marker profiles for all individuals. The elements in *m* are coded as 0, 1 or 2, which is the number of minor alleles at the SNP. ***g*** is an *n*-dimensional random vector representing the genetic background effects. We assume that 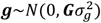 where 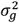 is the genetic variance component, ***G*** is the VanRaden genomic relationship matrix (VanRaden et al., 2008). ***e*** is the residual term and 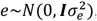 where 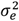 is the residual variance component and ***I*** is the *n* × *n* identity matrix. After solving the linear mixed model, the marker effect was tested using the Wald test statistic *W* = *â*^2^/var(*â*), which approximately follows a *χ*^2^-distribution with one degree of freedom.

Strictly, the model needs to be fitted once for each marker to get the precise test statistic for each marker. But to reduce the computational load, we implemented a commonly used approximate approach, namely the “population parameters previously determined” (P3D) method (Zhang et al., 2010). That is, we only fit the model once without any marker effect (the so-called “null model”), and then we fixed the estimated the variance parameters 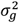 and 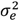 throughout the testing procedure. Then, the test statistic for each marker can be efficiently calculated. GWAS was implemented using R codes developed by ourselves. The variance parameters were estimated by the Bayesian method using the package BGLR (Pérez et al., 2014).

For microbial traits, the significant marker-trait association (MTA) was identified with a threshold of *p* <0.05 after Bonferroni-Holm correction for multiple test (Holm et al. 1979). For fitness phenotypes and alpha-diversity, we used a more liberal threshold of *p* <0.1 after Benjamini-Hochberg correction (Benjamini and Hochberg 1995). For each trait, the proportion of phenotypic variance explained by each MTA (*R*^2^) was calculated as follows: A liner regression model was fitted with all MTAs identified for the trait under consideration. Then, the sum of squares for each MTA as well as the total sum of squares was calculated by ANOVA. The *R*^2^ for each MTA was estimated as the sum of squares of the MTA divided by the total sum of squares.

### GWAS for the presence/absence mode

For microbial traits, we performed in addition a GWAS based on the presence/absence mode (PA-GWAS) in each treatment. Each ASV or taxonomy is considered as present if it is present in more than two replicates (including two). As in the GWAS for abundance, ASVs and taxa with repeatability below 0.1 were filtered out. Those with a presence rate above 95% or below 5% were considered as non-segregated and were also excluded from the analysis. The model for PA-GWAS is a logistic linear mixed model (Chen et al., 2016). Briefly, the model can be described as follows.

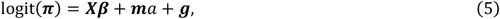

where ***X, β, m***, *a* and ***g*** are the same as in (6). *π* is the vector of conditional probabilities given the covariates, marker effects and the genetic background effects. More precisely, for the *i*-th individual, *π*_*i*_ = *p*(*y*_*i*_ = 1|*X*_*i*_, *m*_*i*_, *g* _*i*_), where *y*_*i*_ is the binary variable indicating the presence (*y*_*i*_ = 1) and absence (*y*_*i*_ = 0), *X*_*i*_ is the *i*-th row of the matrix *X, m*_*i*_ is the *i*-th entry of the vector *m* and *g* _*i*_ is the *i*-th component of the random vector ***g***. The logit function is defined as logit(*x*) = ln(*x*/(1 − *x*)).

Similar to the P3D approach, a null logistic linear mixed model logit(*π*_0_) = ***Xβ* + *g*** was fitted using the penalized quasi-likelihood method (Breslow and Clayton, 1993). The estimated variance components were then fixed throughout the test procedure. A score test was applied to assess the significance of the marker effects.

The PA-GWAS was conducted using the R package GMMAT (Chen et al., 2016).

### Prediction for microbial traits using the genomic data and environmental descriptors

To see the correlation between host genetics and microbiome assemblage, Mantel test was first performed between Rogers’ genetic distance matrix and microbial composition distance matrix only for landraces. After removing the treatment effect using linear model for normalized microbial abundances, the mean value of the residual for each genotype was used to calculate the Euclidean distance. Spearman correlation method was used in mantel function in R. Permutations = 9999.

Next, we investigated the prediction abilities for all microbial traits within each treatment using both the genomic data and the environmental characters. The following three models were implemented. To eliminate the noise of subpopulation effects, we only used the 97 landraces for this part of analysis.

#### Model 1 (genomic prediction)

We applied the genomic best linear unbiased prediction (GBLUP) (VanRaden, 2008) which is the most commonly used model in genomic prediction. The model can be described as follows.

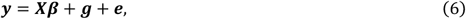

where the notations are the same as in (4). Note that by the use of the VanRaden genomic relationship matrix as the covariance matrix of ***g***, it implicitly modeled the additive effects of all markers.

#### Model 2 (prediction purely based on the environmental characters)

In this model, the genetic effects were replaced by the effects of the environmental characters, which were modeled in a similar way to the GBLUP. More precisely, the model has the following form:

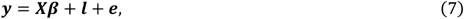

where ***l*** is the *n*-dimensional random vector representing the E-determined values for all individuals. We assume that 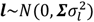 where 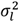 is the corresponding variance component, ***Σ*** is a covariance matrix. Assuming that ***L*** is the *n* × *s* matrix of standardized environmental character records (*s* is the number of environmental characters), we have ***Σ*** = ***LL***^***’***^/*c* where *c* is the mean of all diagonal elements in the matrix ***LL***^***’***^.

#### Model 3 (prediction based on both genomics and environmental characters)

In this approach, we combined the genomic data and the Es in a multi-kernel model, which is of the following form:

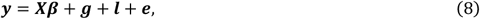

where the notations were inherited from (6) and (7).

The prediction abilities of the above three models were assessed in a leave-one-out cross-validation scenario. That is, each individual was predicted once using a training set consisting of all other individuals. Thus, for each trait the prediction model was fitted *n* times. After we obtained the predicted values of all individuals, the prediction ability was calculated as the correlation between the predicted and observed values. The standard error was estimated using the bootstrap approach (Efron, 1979).

All prediction models were implemented using the R package BGLR (Pérez et al., 2014) and rrBLUP (Carley et al., 2017).

### Prediction for plant phenotypes using the genomic and microbiome data

We explored the possibility of predicting the three fitness phenotypes and ionome traits in each treatment using the genomic data and microbiomes. As in the last subsection, we focused on the subpopulation of 97 landraces.

#### Scenario 1 (prediction based on microbiomes only)

In this scenario, we considered 9 cases, in which the phenotypes were predicted using bacteria in the root sample (BA_RO), in the rhizosphere sample (BA_RH), fungi in the root sample (FU_RO), in the rhizosphere sample (FU_RH), bacteria in both samples (BA), fungi in both samples (FU), both types of microbiomes in the root sample (RO), in the rhizosphere sample (RH), and both types of microbiomes in both samples (ALL). The model can be uniformly described as follows:

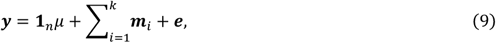

where ***m***_*i*_ is an *n*-dimensional trait values for all individuals determined by a certain type of microbiome in a specific sample, *k* can be 1 (BA_RO, BA_RH, FU_RO, FU_RH), 2 (BA, FU, RO, RH), or 4 (ALL), other notations are the same as in (8). We assume that 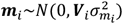, where 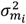 is the corresponding variance component, ***V***_*i*_ is a covariance matrix derived from the microbiome ASVs. Assuming that ***M***_*i*_ is the *n* × *t* matrix of standardized records of microbiome ASVs (t is the number of different ASVs), we have ***V***_*i*_ = ***M***_*i*_***M***_*i*_^***’***^/*c*_*i*_ where *c*_*i*_ is the mean of all diagonal elements in the matrix ***M***_*i*_***M***_*i*_^***’***^.

#### Scenario 2 (prediction based on both microbiomes and genomic data)

In this scenario, the 9 cases in Scenario 1 were combined with genomic data (G_BA_RO, G_BA_RH, G_FU_RO, G_FU_RH, G_BA, G_FU, G_RO, G_RH, G_ALL). The models are of the following form:

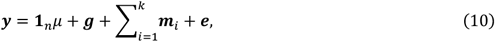

where the notations were adopted from (8) and (11).

As in the last subsection, the prediction abilities were evaluated in a leave-one-out cross-validation scenario. Prediction models were implemented using the R package BGLR.

### Effects of source environmental factors on specific microbial assemblies

To explore the environmental legacy of native habitats in relation to specific microbial variations among landraces, we performed network analyses of rhizosphere and root microbial indicators. We then aimed to understand the connections between bacterial and fungal taxa intimately associated with the microbiome of roots and rhizospheres. To this end, we used the function “multipatt” in the R package indicspecies (De Cáceres et al., 2020) to identify those microbial phylotypes that were significant indicators of microbial zASVs roots and rhizosphere (i.e., roots, rhizosphere or roots + rhizosphere) compared with bulk soil. We then conducted a correlation network conformed by taxa associated with the root and rhizosphere microbiomes. We calculated all pairwise Spearman correlation coefficients among these microbial taxa and kept all positive correlations. We further identified microbial modules (clusters of taxa highly correlated with each other) using Gephi (https://gephi.org/). We determined the proportion of modules by calculating the standardized (0-1) average of all taxa within each module, so that all taxa equally contribute to each module. This information was then correlated (Spearman) with environmental conditions. Mean annual temperature and precipitation were obtained from the WorldClim database (https://www.worldclim.org/). Other environmental descriptors were determined as explained above. Structural equation modelling (SEM) was conducted to provide a system-level understanding on the direct and indirect associations between environmental factors, the proportion of modules and that of selected taxa from above-explained analyses. Because some of the variables introduced were not normally distributed, we used bootstrap tests in these SEMs. We evaluated the fit of these models using the model χ^2^-test, the root mean squared error of approximation and the Bollen– Stine bootstrap test.

### Environmentally adaptive loci and microbiome relatedness across abiotic stresses

To determine if the environmentally associated loci are contributing to microbiome adaptation to abiotic stresses, we used a representative set of natural varieties e.g. 97 landraces accessions covering typical geographical range. Prior to analysis, PCA was conducted based on the BLUEs for each treatment and compartment to extract major sources of variance from bacterial and fungal microbial community data. The first five PCs were obtained for downstream analyses. PCA was performed using the prcomp function in R. In addition, we selected 18 individual ASVs belonging to *Oxalobacteraceae* to be predicted by Random Forest models. To improve model accuracy, feature selection was conducted prior to model building to eliminate unimportant or redundant environmental variables by identifying those with significant associations to an outcome variable. The feature selection method Boruta was employed to identify environmental aspects that describe significant variation in the PCs and ASVs using Boruta::boruta() (v7.0.0) (Kursa and Rudnicki, 2010).

The subset of boruta-identified environmental variables (Supplementary Dataset 12) for each ASV were used for Random Forest model construction. This model works under the expectation that a response variable can be described by several explanatory variables through the construction of decision trees. Thus, each Random Forest model is representative of the non-linear, unique combination of explanatory variables that describe variation in a response variable. Random Forest models were built using RandomForest::randomForest() function under default parameters, 5000 trees were built and one third of the number of explanatory variables were tried at each split (Liaw and Wiener et al., 2002). Random Forest models were trained with 80% of the data and validated with the remaining 20% test set. Model success was evaluated with percent error explained, Nash-Sutcliffe efficiency (NSE), mean absolute error (MAE), and mean squared error (MSE). Using constructed Random Forest models, ASVs were predicted for 1,781 genotyped landraces in Mexico. These landraces were genotyped as a part of the Seeds of Discovery project (SeeD).

We conducted genome wide association studies (GWAS) to measure the associations between SNPs of landrace genotypes and predicted microbial traits, as well as the associations between SNPs and the environmental variables used to predict the microbial traits. SNPs were filtered for minor allele frequency >1%. We applied the method as previously described (Gates et al., 2019), using a linear model to fit the genotypic data and each microbial trait and environmental variable for Mexican landrace accessions. The first five eigenvectors of the genetic relationship matrix were included in the model to control for population structure. To control for the number of false positive tests, we re-calibrated the *p*-values using the false discovery rate (FDR) control algorithm (François et al., 2016) and selected significant SNPs based on the calibrated results. To test if GWA hits based on the prediction is significantly better in capturing top GWA hits of observed data than random, we conducted a permutation test and compared the median p-value of GWA hits of observed data that are around 200kb of the top 100 prediction-based GWA hits and the median p-value of random selected GWA hits based on 10000 permutations.

### Association of allele frequency with soil nitrogen and microbial taxa

To identify whether the microbiome is associated with environment and maize phenotypes, we performed allelic variation analysis of Zm00001d048945 using an SNP dataset of CIMMYT landraces accessions obtained from a previous study (Navarro et al., 2017). We extracted the genotypic information of top SNPs of the target gene Zm00001d048945 for all tested landraces. We divided maize landraces into 20 groups based on the total soil nitrogen content (%) of their sampling sites (Shangguan et al. 2014). We calculated the mean total nitrogen, the minor allele frequencies (MAF) of the target SNPs, and the mean predicted ASV abundance for each group of landraces. Pearson correlation was conducted to test the correlations between MAF and total nitrogen content, and between MAF and ASV abundance.

### Candidate gene validation by independent transposon insertion alleles

Gene expression for Zm00001d048945 was explored in qTeller (https://qteller.maizegdb.org/), which allows to compare gene expression across different tissues from multiple data sources. Gene expression data was extracted from different organs (seed, root, tassel/silk, internodes and leaf) and specific tissues such as the root meristematic zone, elongation zone, stele and cortex. The gene encoded protein annotation was inferred from UniProt database (https://www.uniprot.org/). We next identified potential loss-of-function mutations by exploring the sequence indexed collection BonnMu (Marcon et al., 2020). Induced maize mutants of the BonnMu resource derive from Mutator-tagged F_2_-families in various genetic backgrounds, such as B73 and F7. We identified two insertion lines, BonnMu-8-D-0170 (B73) and BonnMu-F7-2-F-0598 (F7), harboring insertions 1,264 bp upstream of the start codon ATG and in the second exon of Zm00001d048945, respectively. These two families were phenotyped in paper-roll culture (Yu et al., 2021) and the seedling plants were scanned using the scanner Expression 12000XL (Epson, Suwa, Japan). Lateral roots were counted and the density was normalized with the measure number of lateral roots per cm length of primary root. Statistical analyses were performed by pair-wise Students *t* test with *F* statistics.

### Association of relative abundance of *Massilia* with lateral root density

To understand the relationship between *Massilia* and the formation of lateral roots, root system architecture and morphology of 129 maize genotypes was scanned with an Epson Expression 12000XL scanner. Lateral root density was determined by manual calculation as the number of emerged lateral roots per length (cm) of the main root. The linear correlation was plotted between lateral root density and relative abundance data of *Massilia* ASVs using R (v4.1.0).

### Data availability

All raw maize genotyping data, bacterial 16S and fungal ITS data in this paper were deposited in the Sequence Read Archive (http://www.ncbi.nlm.nih.gov/sra) under the BioProject ID PRJNA889703. The SSUrRNA database from SILVA database (release 138, 2020, https://www.arb-silva.de/) and UNITE database (v8.3, 2021, https://unite.ut.ee/) were used for analysing the bacterial 16S and fungal ITS sequences, respectively. We deposited customized scripts in the following GitHub repository: https://github.com/Danning16/MaizeMicrobiome2022. All statistical data are provided with this paper.

## Acknowledgement

We thank Candice Gardner (United States Department of Agriculture, Ames, US) and the International Maize and Wheat Improvement Center (CIMMYT) for germplasm contribution. We thank Angelika Glogau, for soil and plant nutrient determination and Selina Siemens and Alexa Brox for soil and root DNA extractions (University of Bonn, Bonn, Germany). We thank Yayu Wang and Huan Liu (State Key Laboratory of Agricultural Genomics, BGI-Shenzhen, Shenzhen, China) for providing us the SNP matrix data in foxtail millet. We thank Daliang Ning and Jizhong Zhou (University of Oklahoma, Norman, USA) for suggestions on the microbiome data analysis. This work is supported by Deutsche Forschungsgemeinschaft (DFG) grants HO2249/9-3, HO2249/12-1 to F.H. and YU272/1-1 and Emmy Noether Programme 444755415 to P.Y., the German Excellence Strategy – EXC 2070 – grant 390732324 to P.Y. and G.S., the Bundesministerium für Bildung und Forschung (BMBF) grant 031B195C to F.H. and DFG Priority Program (SPP2089) “Rhizosphere Spatiotemporal Organisation - a Key to Rhizosphere Functions” grant 403671039 to F.H. and P.Y. X.C.’s research is supported by The Changjiang Scholarship, Ministry of Education, China, State Cultivation Base of Eco-agriculture for Southwest Mountainous Land (Southwest University, Chongqing, China), and the National Maize Production System in China (grant no. CARS-02-15).

