## Supplemental figures and tables for "Heritable microbiome variation is correlated with source environment in locally adapted maize varieties" (from another version of this preprint: version 1, 2023-01-10)

**Supplementary Figures and Tables**

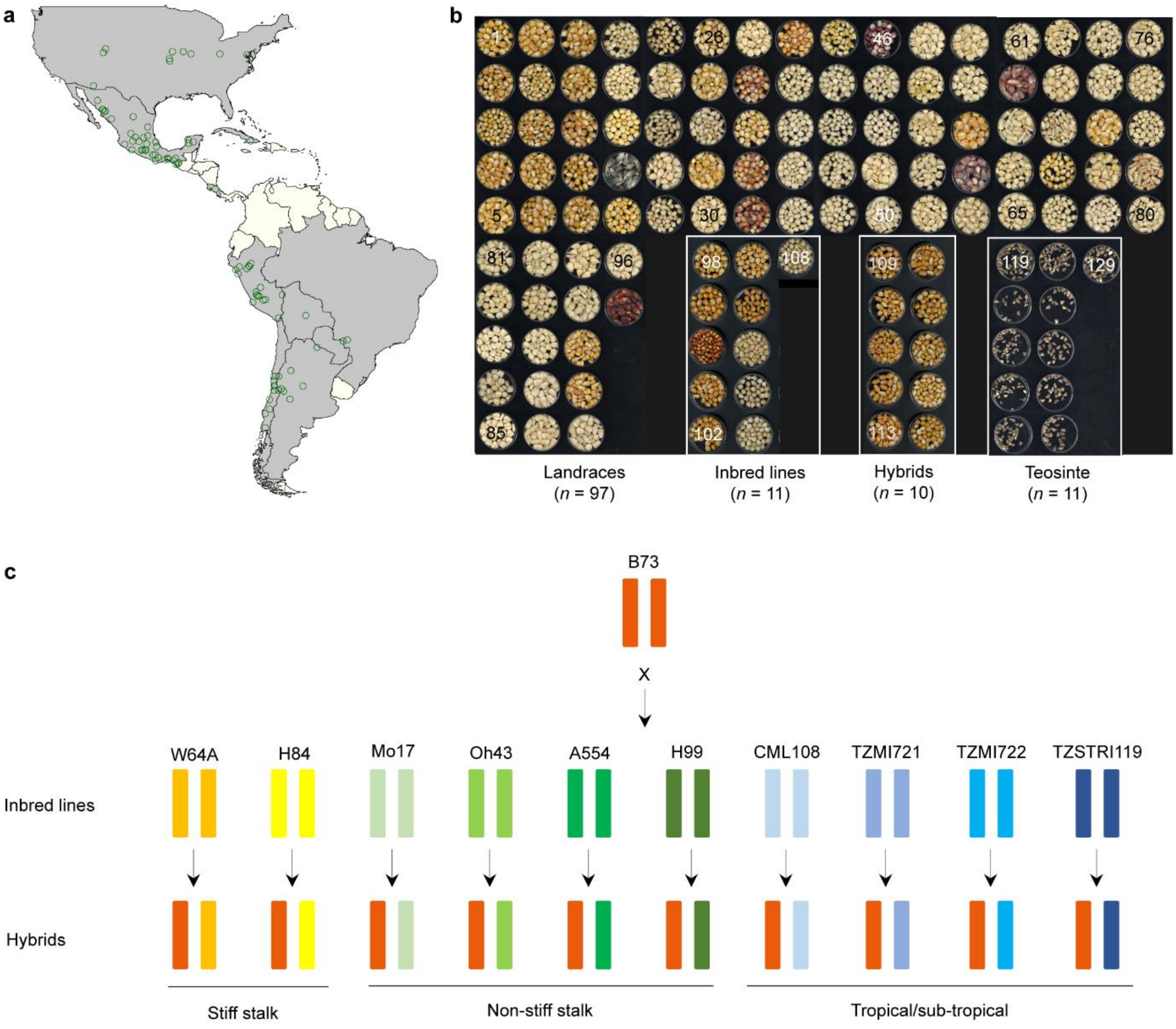


**Supplementary Figure 1. Germplasm used in this study. a**, Geographical coordinates of the 97 collected maize landraces are highlighted by green circles. **b**, Seed phenotypes of 97 landraces, 11 teosinte accessions, 11 inbred lines and 10 hybrids. **c**, Design of crosses of 10 inbred lines with the recurrent mother plant B73 to generate hybrids. The parental lines can be divided into three sub-populations (stiff stalk, non-stiff stalk, tropical/sub-tropical).

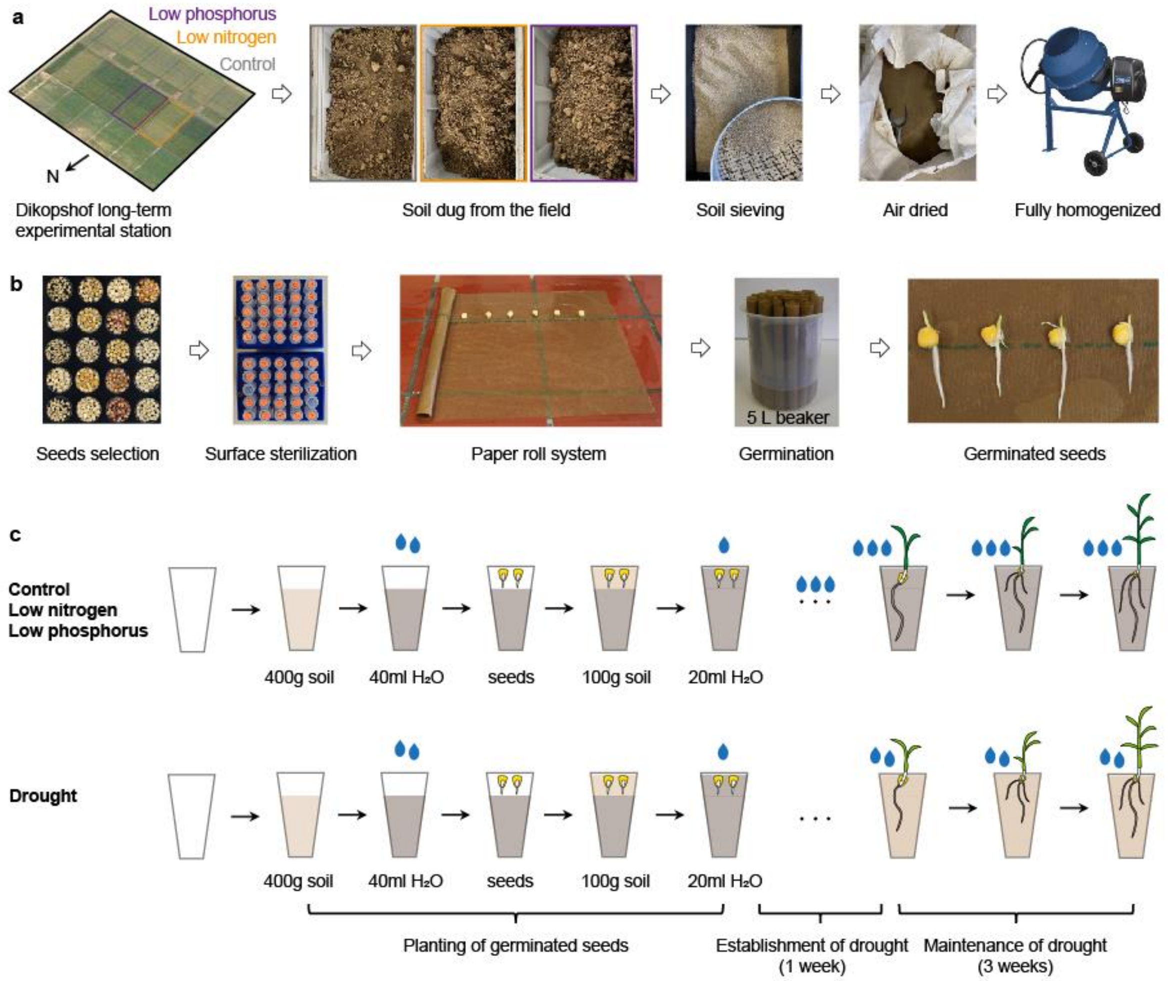


**Supplementary Figure 2. Soil pot experiments. a**, Natural soils with different nutrient levels were dug from Dikopshof long-term experimental station (see methods). The soils were then sieved with a 4 mm sieve, air-dried and homogenized. **b**, Seed sterilization and germination. Only seeds with similar phenotypes were sterilized and germinated in a paper-roll system until the primary root had a length of 1-2 cm. **c**, Transfer to the soil-filled pots. The germinated seeds were transferred to the soil-filled pots with sterilized water. Drought was established by suspending watering for one week and maintenance of low soil water moisture at mild stress level described in Supplementary Figure 33. The plants were grown in the soil-filled pots for 4 weeks.

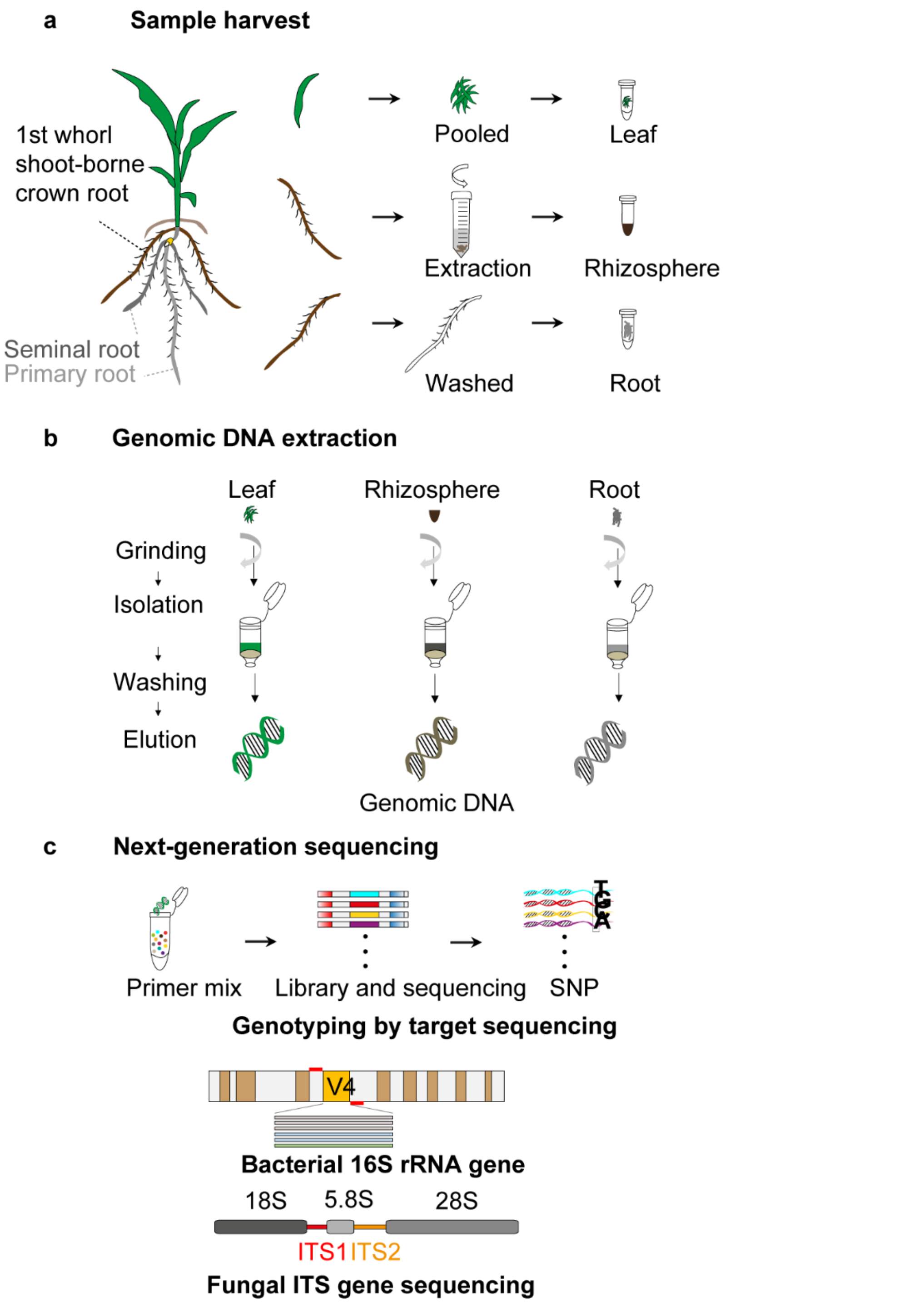


**Supplementary Figure 3. Sample harvest and sequencing strategy. a**, Harvesting of shoot, rhizosphere and root samples. Fresh leaf tip samples were dissected and pooled from several maize plants grown under different treatment conditions. Rhizosphere sample was extracted from the 1$^{st}$ shoot-borne crown root with tightly attached soil. The same whorl of shoot-borne roots was separately dissected and immediately washed, dried with clean tissue, followed by flash freezing in liquid nitrogen. **b**, Extraction of genomic DNA from leaf, rhizosphere and root samples. **c**, Next-generation sequencing. The genomic DNA from pooled leaf samples was genotyped by genotyping by target sequencing (see Methods). The genomic DNA from rhizosphere and root samples was sequenced by 16 rRNA (V4 region) gene and ITS1 gene sequencing for bacterial and fungal community analyses.

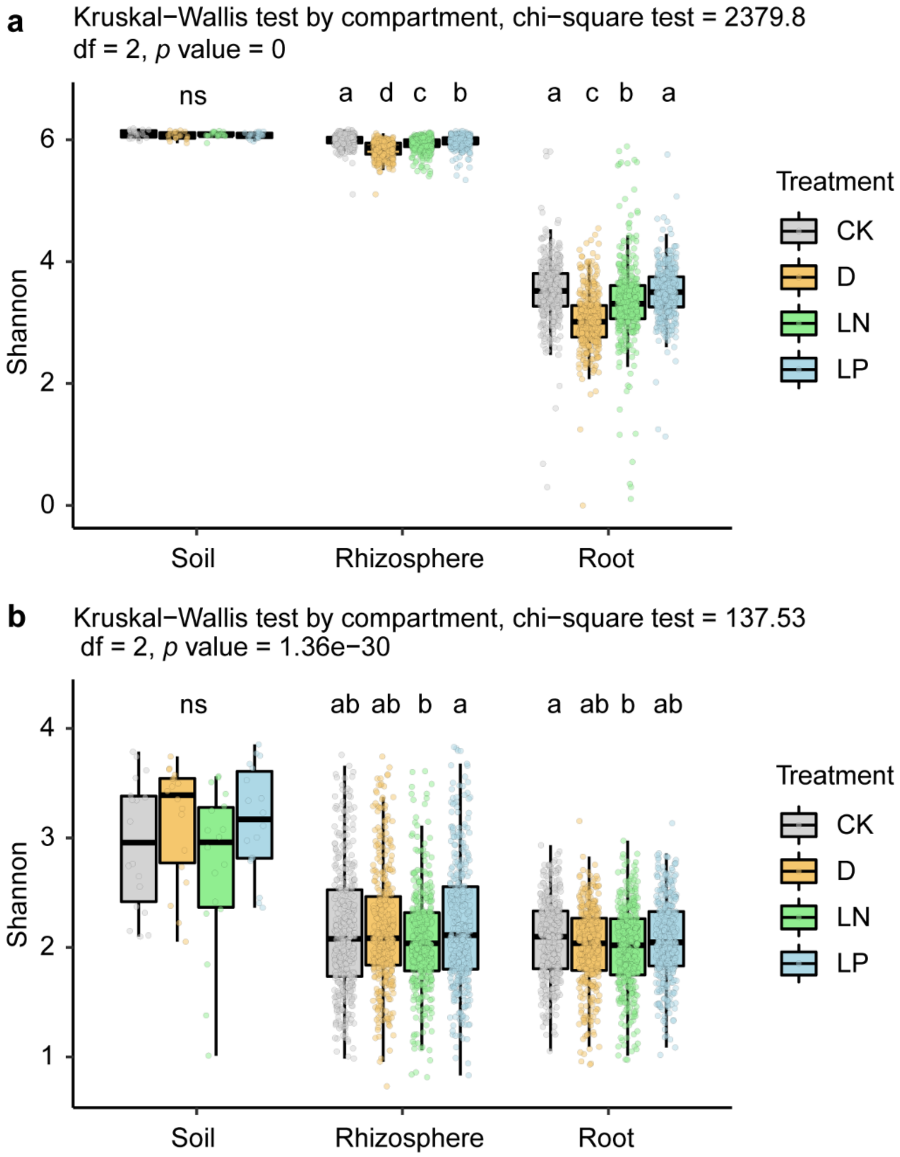


**Supplementary Figure 4. Microbial diversity across treatments in different compartments.** Changes in bacterial (**a**) and fungal (**b**) alpha-diversity estimated by Shannon's diversity index. Compartment significances were calculated using a two-tailed Kruskal-Wallis test with post-hoc Wilcoxon (Benjamini-Hochberg adjusted). Different letters indicate significance between different treatments ($p$ <0.05). CK, control; D, drought; LN, low nitrogen; LP, low phosphorus. ns, not significant. Box plots include the median (the horizontal line inside the boxes), 95% confidence intervals (the black vertical lines) and the distribution of frequency (width of the differently colored zones). For **a** and **b**, $n$ = 18 for soil under each treatment, $n$ = 387 for rhizosphere and root under each treatment. All samples are biologically independent.

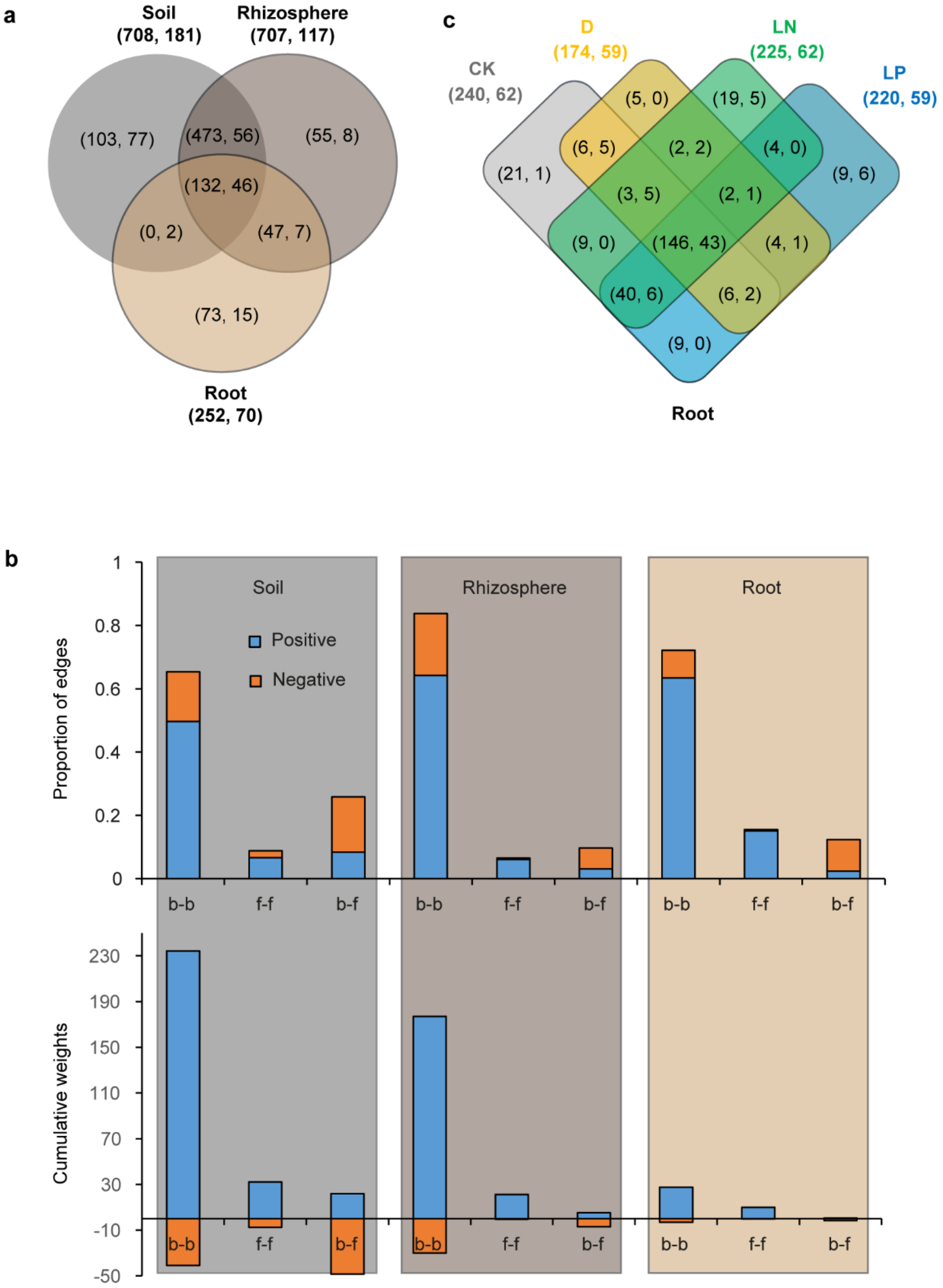


**Supplementary Figure 5. Intra- and inter-kingdom network analysis. a**, 3-way Venn diagram indicates the overlapping patterns of bacterial and fungal interacting ASVs across the compartments. The first and second number in the brackets indicate the bacterial and fungal ASVs respectively. **b**, Bar plot indicates the network associations measured by proportion of edges and cumulative weights within and between different microbial kingdoms. b-b, bacterial-bacterial interactions; f-f, fungal-fungal interactions; b-f, bacterial-fungal interactions. **c**, 4-way Venn diagram indicating the overlapping patterns of bacterial and fungal interacting ASVs across treatments in the root. The first and second number in the bracket indicate the bacterial and fungal ASVs respectively.

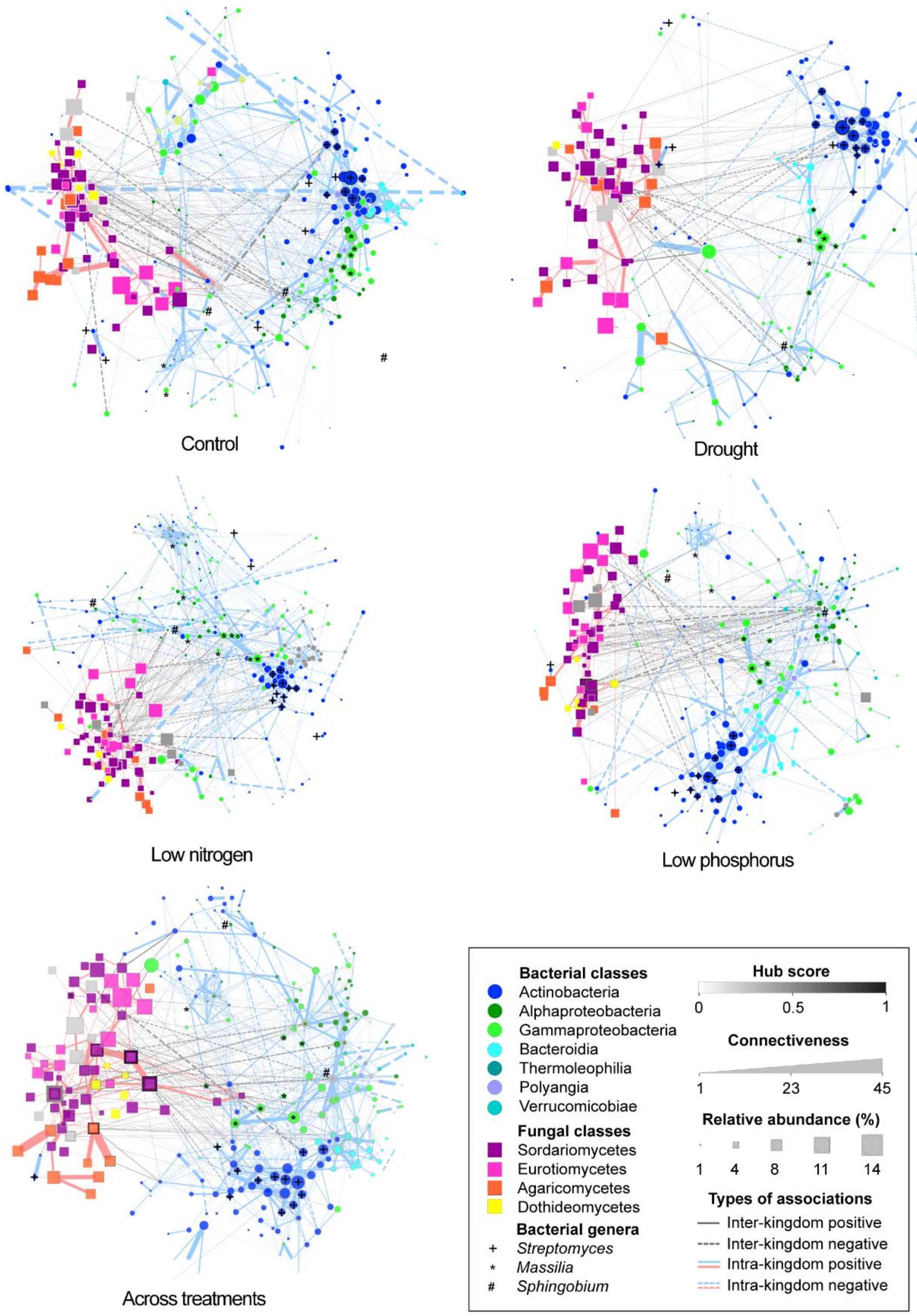


**Supplementary Figure 6. Intra- and inter-kingdom root network associations under different stress treatments.** A sparse inverse covariance estimation for ecological association inference (SPIEC-EASI) based network association analysis of root-associated microbial ASVs (RA >0.05%; Prevalence >10%) across different treatments. Each node corresponds to an ASV and was color coded according to the family taxa. The size of nodes reflects the relative abundance in the root. Edges between nodes correspond to either positive (solid lines) or negative (dotted lines) associations and color codes of edges indicate inter- or intra-kingdom associations between ASVs. Connectiveness between nodes corresponds to the degree of associations. Keystone ASVs in microbial networks were reflected by hub scores corresponding to the thickness of the nodes.

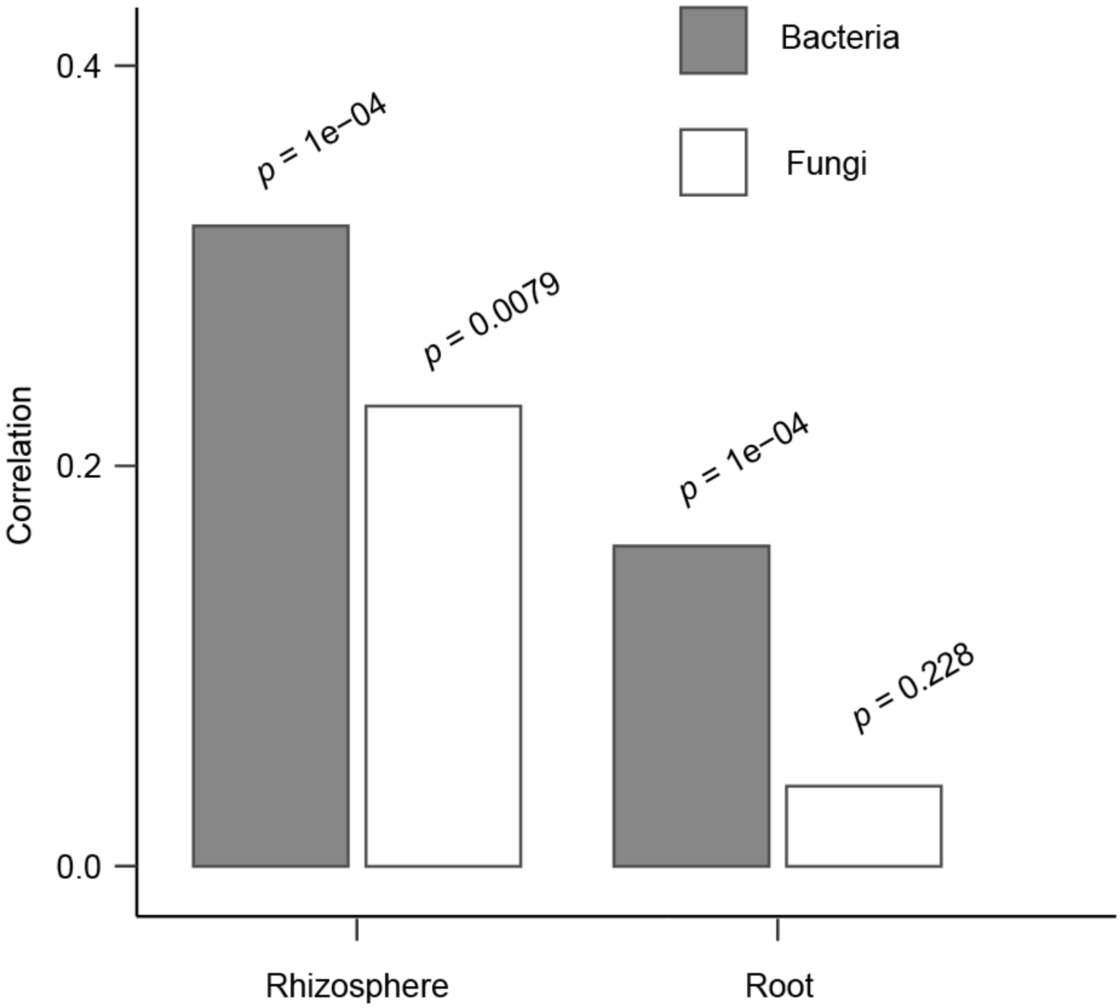


**Supplementary Figure 7. Mantel's statistic tests for correlation between host genotypic distance and microbiome distance.** The exact *p* values are indicated.

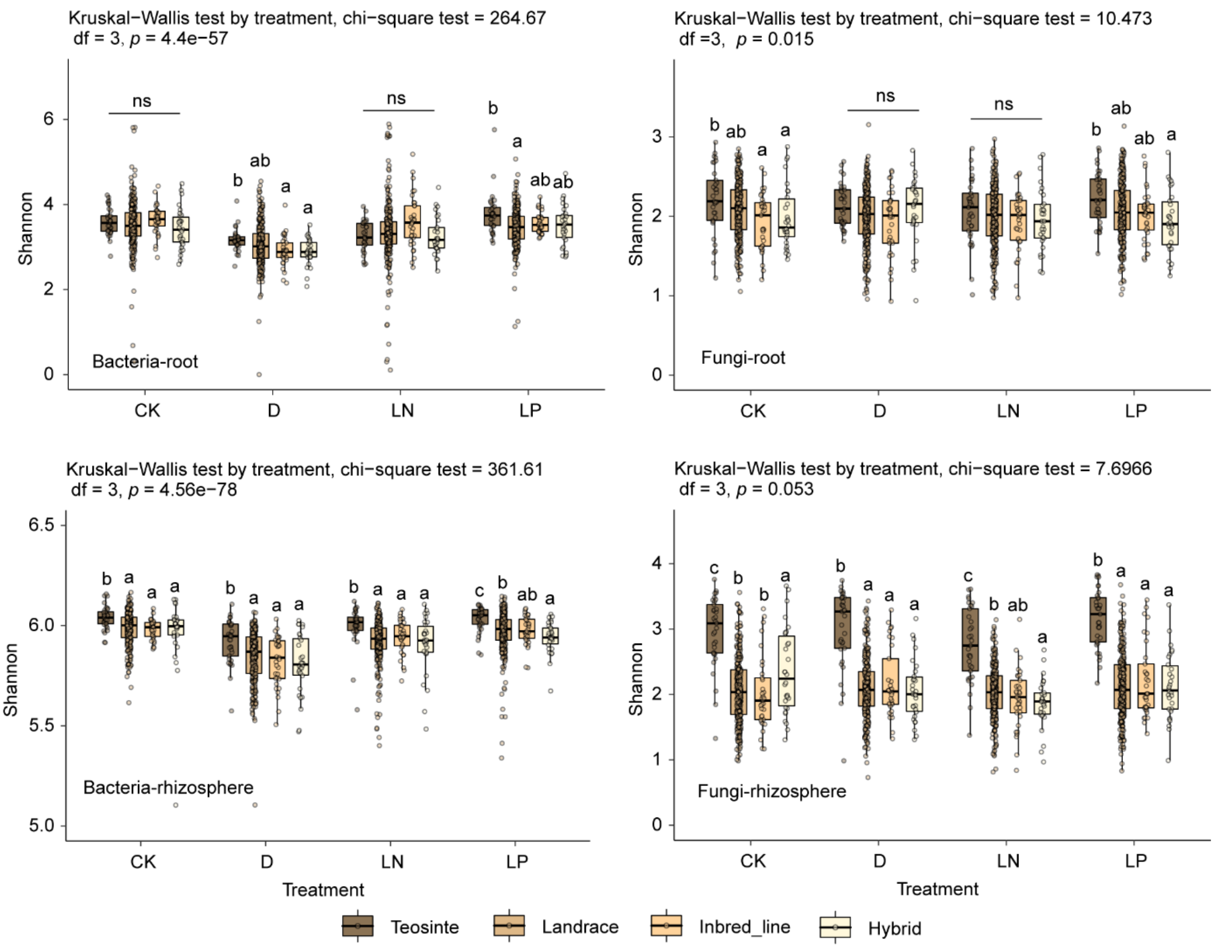


**Supplementary Figure 8. Microbial diversity across germplasm groups in different treatments.** Changes in bacterial and fungal alpha-diversity estimated by Shannon's diversity index in the root and rhizosphere. Differences among treatment were calculated using a two-tailed Kruskal-Wallis test with post-hoc Wilcoxon test (Benjamini-Hochberg adjusted). Different letters indicate significance among germplasm groups ($p$ <0.05). CK, control; D, drought; LN, low nitrogen; LP, low phosphorus. ns, not significant. Box plots include the median (the horizontal line inside the boxes), 95% confidence intervals (the black vertical lines) and the distribution of frequency (width of the differently colored zones). $n$ = 33 for teosinte, $n$ = 291 for landraces, $n$ = 33 for inbred lines, $n$ = 30 for hybrids under each treatment. All samples are biologically independent.

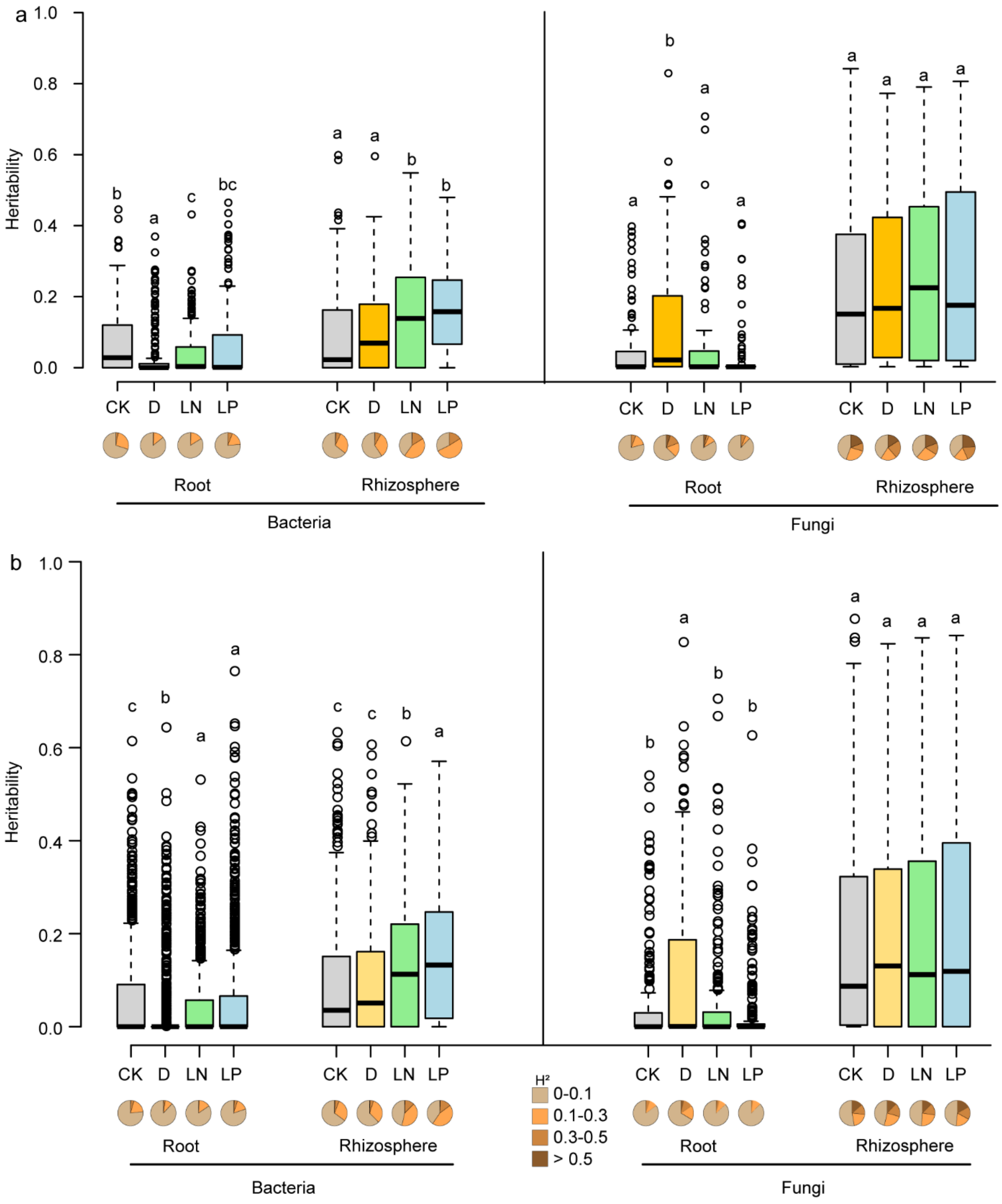


**Supplementary Figure 9. Heritability estimation of microbial traits. a, Heritability of all microbial** genera (**a**) and ASVs (**b**) under four treatments for both bacteria and fungi. The broad-sense heritability ($H^2$) was calculated using highly abundant bacterial ($n$ = 815) and fungal ($n$ = 248) ASVs across all samples. Significant differences are indicated for each compartment among treatment groups at $p$ <0.05 after Benjamini-Hochberg adjustment (Kruskal-Wallis test, Dunn's *post-hoc* test). Boxes span from the first to the third quartiles, centre lines represent the median values and whiskers show data lying within the 1.5× interquartile range of the lower and upper quartiles. Data points at the ends of whiskers represent outliers. The pie chats indicate the proportional distribution of heritability frequencies. CK, control; D, drought; LN, low nitrogen; LP, low phosphorus.

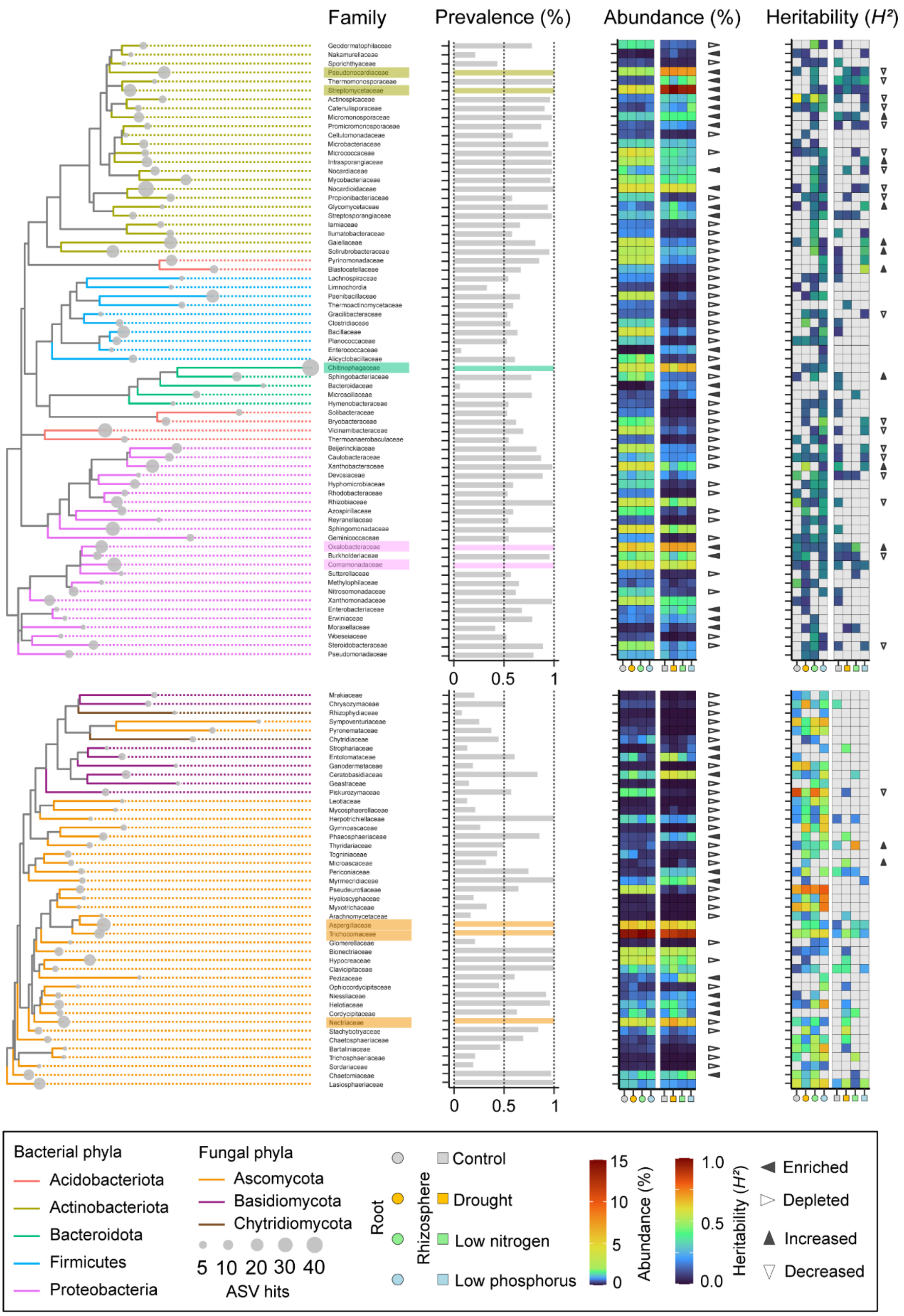


**Supplementary Figure 10. Maximum-likelihood phylogeny of dominant microbial families (n > 5).** Circle sizes along the branches of the tree indicate the number of ASVs observed in association with microbial families. Colour coded families are clustered at the phylum level. Bar plots describe the prevalence according to the proportional sample size. The heatmaps illustrate the standardized mean relative abundance and the estimated heritability of microbial families from the root to the rhizosphere. Triangles represent the enrichment or depletion of microbial families, and increased or decreased heritability from the root to the rhizosphere. The significance levels were controlled at two levels (*: $p$ <0.05; **: $p$ <0.01).

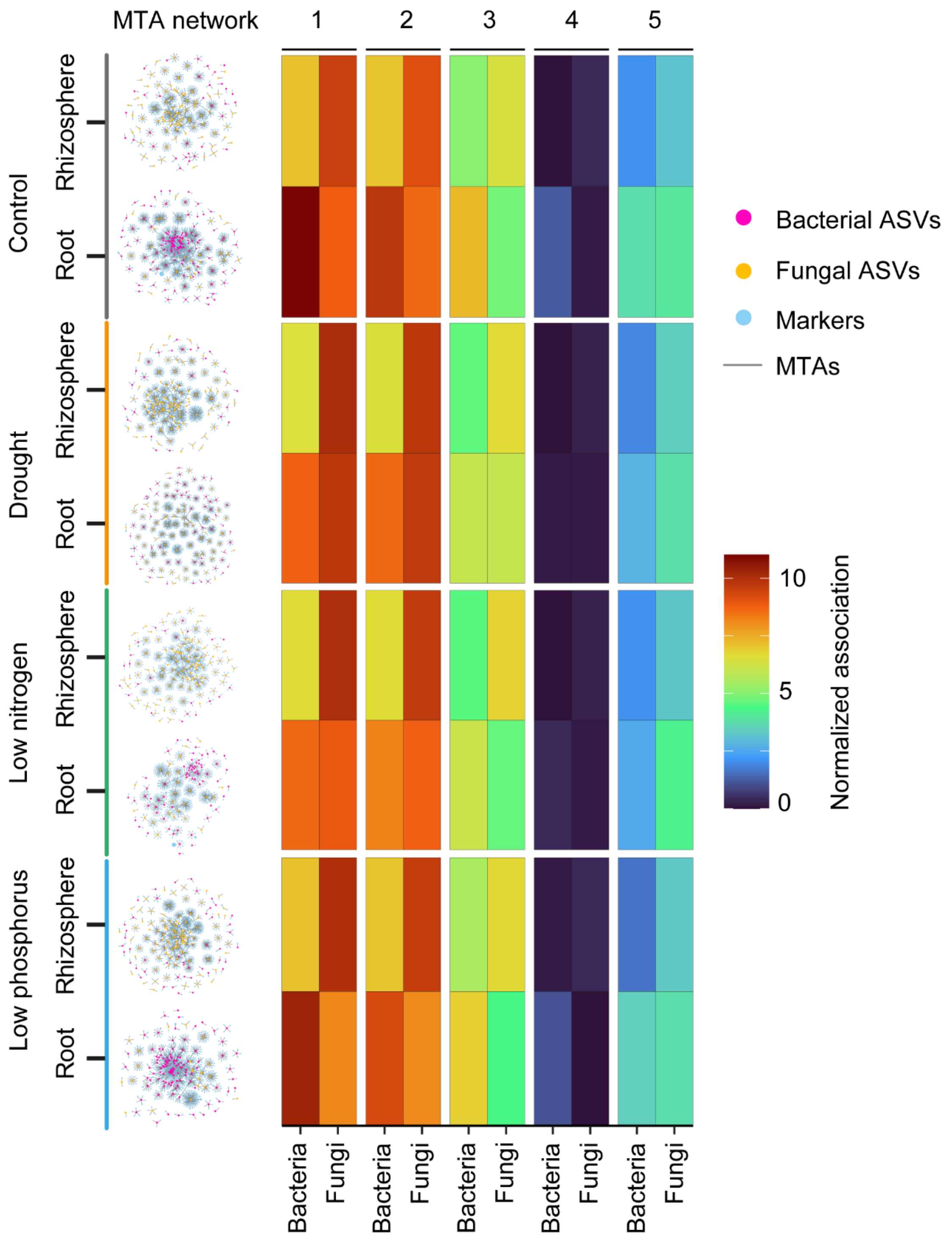


**Supplementary Figure 11. Network visualization of host genetic markers and association with microbial ASVs.** A sparse inverse covariance estimation for ecological association inference (SPIEC-EASI) based network association analysis of root-associated microbial ASVs (RA >0.05%; Prevalence >10%) across different treatments and genotypes. Each node corresponds to an ASV and was color coded according to the microbial classes. The size of nodes reflects the relative abundance in the root. Edges between nodes correspond to either positive (solid lines) or negative (dotted lines) associations and color codes of edges indicate inter- or intra-kingdom associations among ASVs. Connectiveness between nodes corresponds to the degree of associations. Keystone ASVs in microbial networks are reflected by hub score corresponding to the thickness of node. MTA, marker trait association. Normalized associations were illustrated by $\log_2$ transformation for total MTAs (1), unique markers (2), unique ASVs (3), associations per marker (4) and associations per ASV (5).

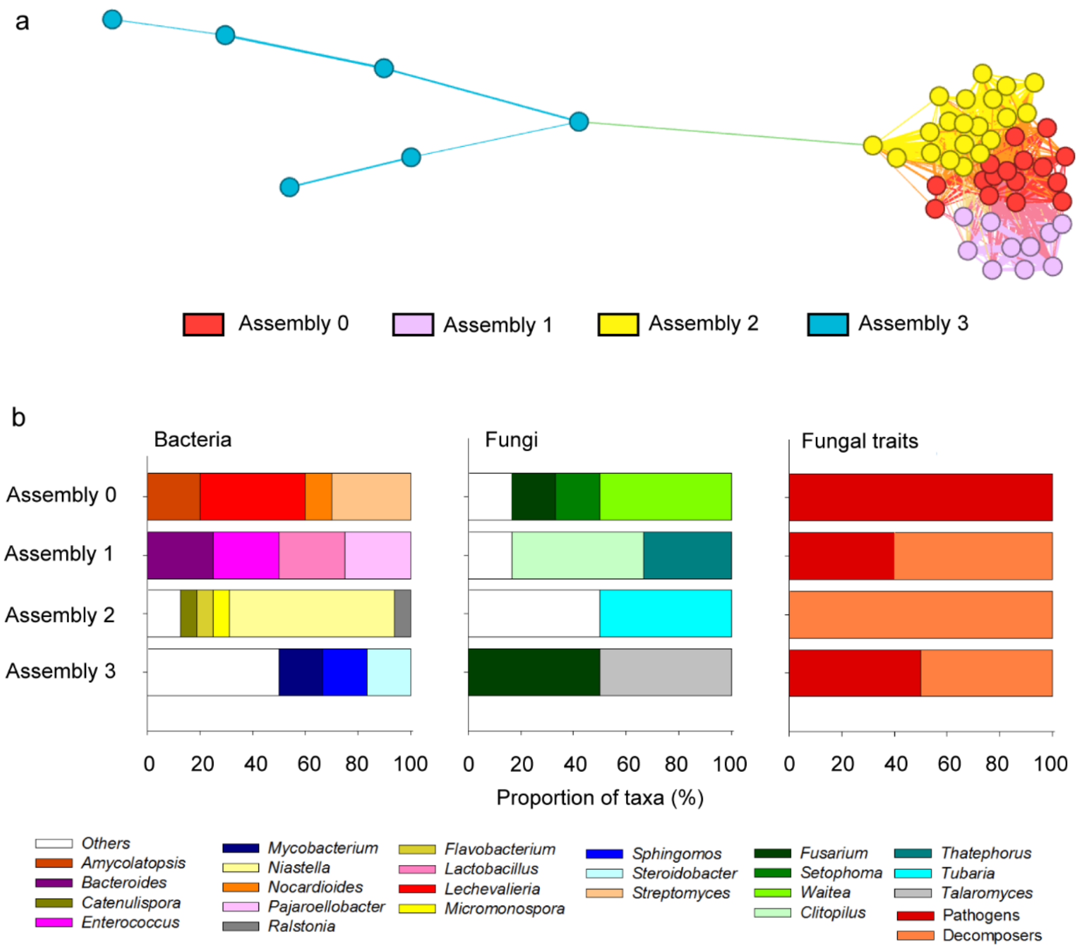


**Supplementary Figure 12. Correlation network including independent assemblies conformed by taxa associated with the root and rhizosphere microbiomes.** Panel A includes a correlation network considering only microbial indicators of rhizosphere and root environments as determined in the Method section. Different assemblies represent modules within the ecological network including microbial taxa highly co-occurring with each other. Panel B represents the main taxa including in each other modules reported in Panel A.

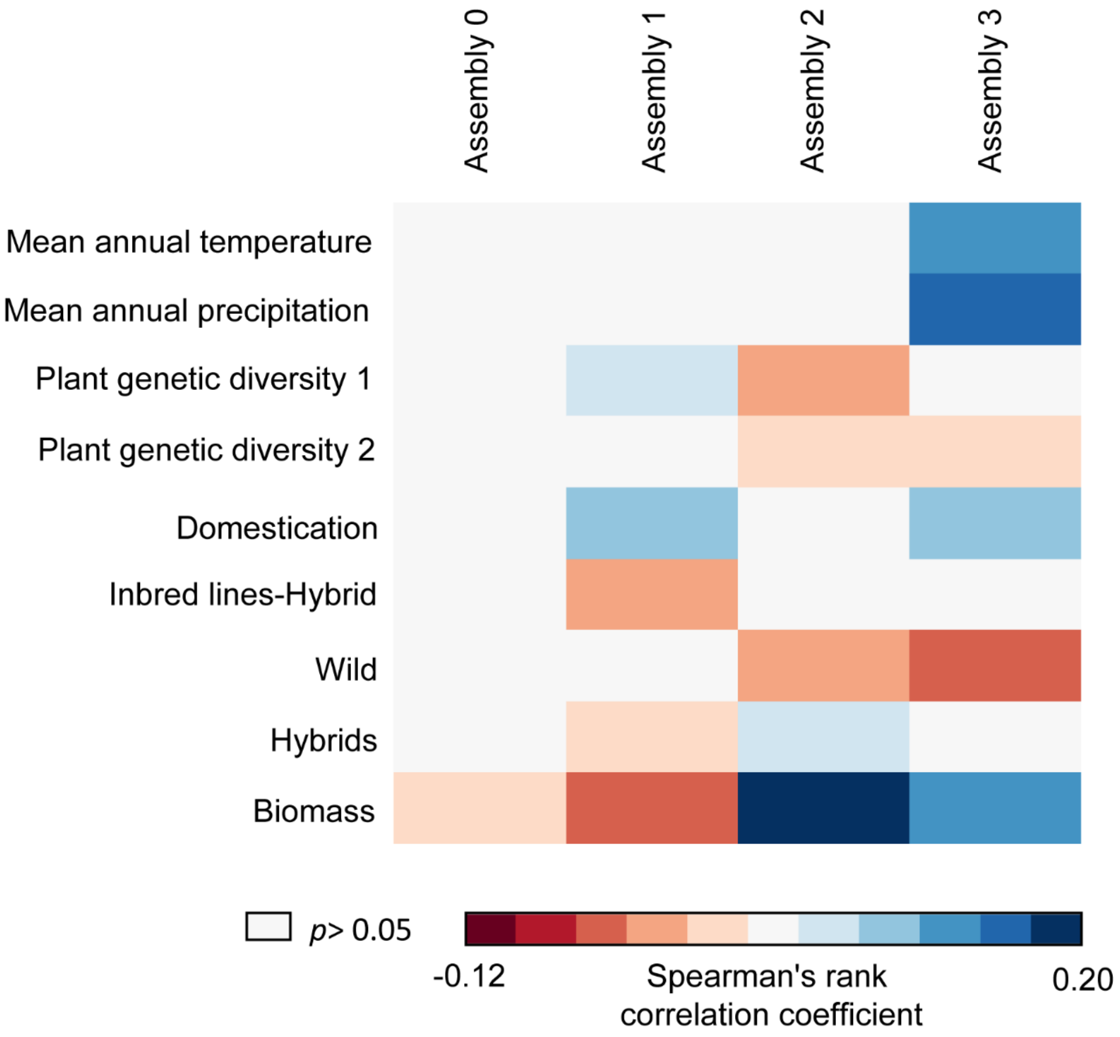


**Supplementary Figure 13. Spearman correlations between legacy and genotype characteristics and the correlation of the four independent microbial assemblies in Supplementary Figure 12.** Plant genetic diversity 1 and 2 are determined as the axes of a NMDS (Non-metric Multidimensional Scaling) analysis including maize genetic variability. Mean annual precipitation and mean annual temperature represent natural precipitation and temperature conditions in the places of origin for maize varieties. They are included in our model as climatic legacies as our microbiomes were determined in growth chamber experiments under control conditions.

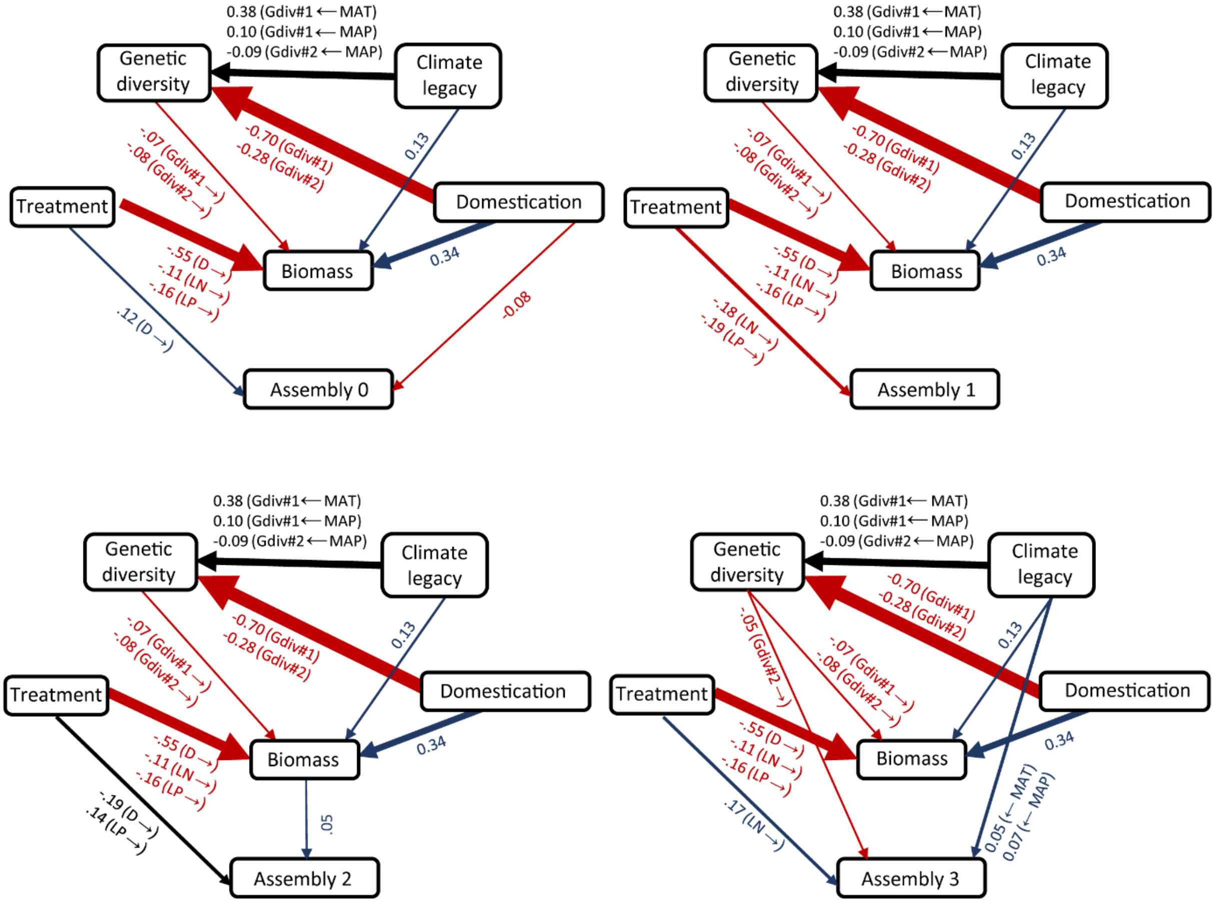


**Supplementary Figure 14. Structural equation modeling exploring the direct and indirect effects of climatic legacies, genotype diversity, treatments, domestication and biomass on the four independent microbial assemblies in Supplementary Figure 12.** Plant genetic diversity 1 (Gdiv#1) and 2 (Gdiv#2) are determined as the axes of a NMDS (Non-metric Multidimensional Scaling) analysis including maize genetic variability. Mean annual precipitation (MAP) and mean annual temperature (MAT) represent field temperature and precipitation conditions in the places of origin for maize varieties. They are included in our model as climatic legacies as our microbiomes were determined in growth chamber experiments under control conditions. CK, control; D, drought; LN, low nitrogen; LP, low phosphorus.

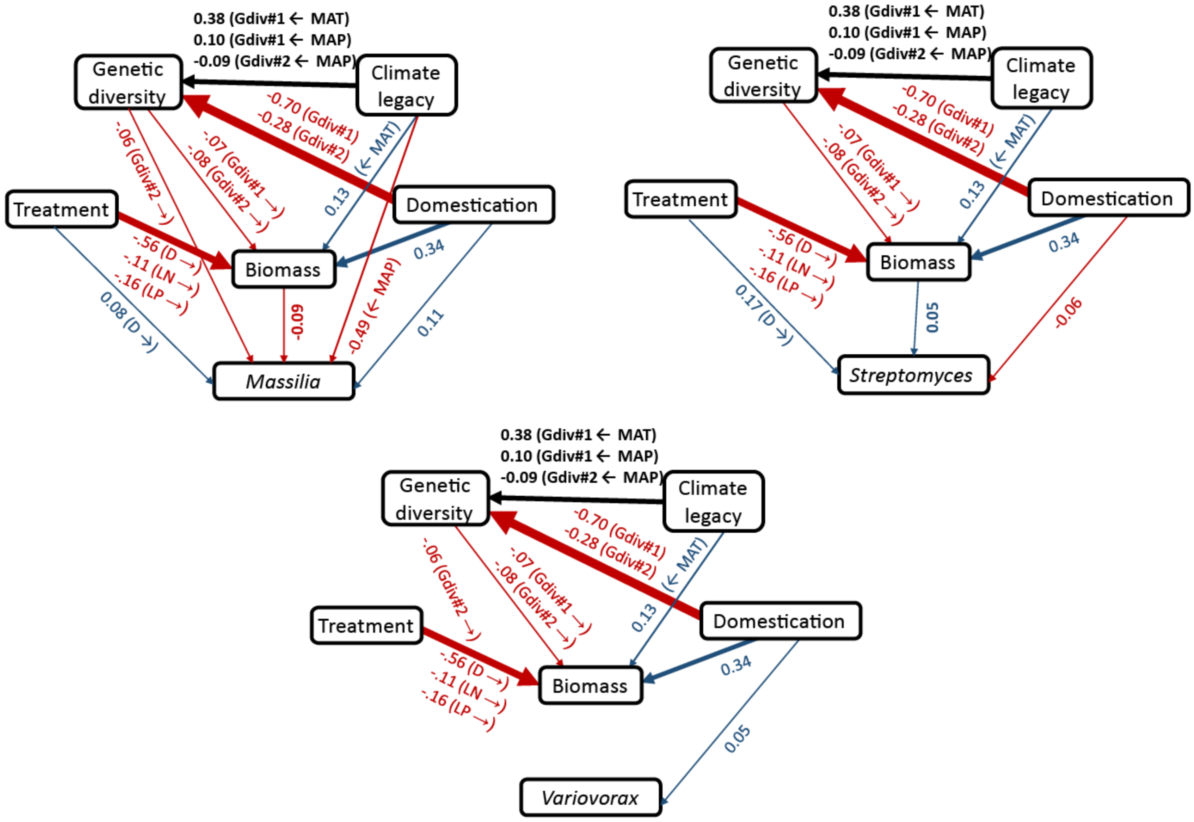


**Supplementary Figure 15. Structural equation modeling exploring the direct and indirect effects of climatic legacies, genotype diversity, treatments, domestication and biomass on keystone bacterial genera.** Plant genetic diversity 1 (Gdiv#1) and 2 (Gdiv#2) are determined as the axes of a NMDS (Non-metric Multidimensional Scaling) analysis including maize genetic variability. Mean annual precipitation (MAP) and mean annual temperature (MAT) represent field temperature and precipitation conditions in the places of origin for maize varieties. They are included in our model as climatic legacies as our microbiomes were determined in growth chamber experiments under control conditions. CK, control; D, drought; LN, low nitrogen; LP, low phosphorus.

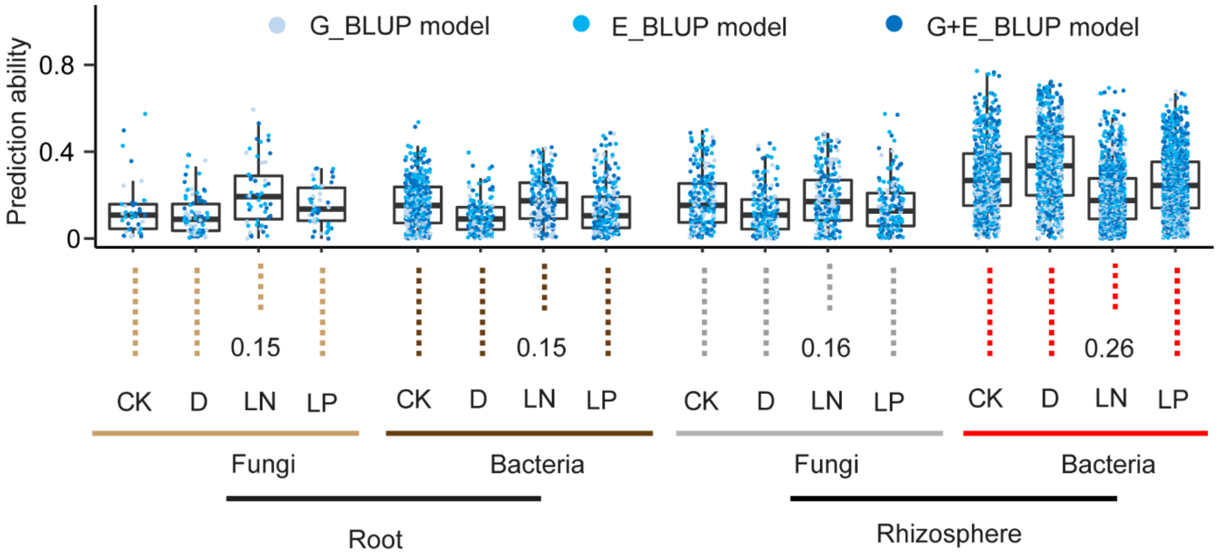


**Supplementary Figure 16. Genomic and environmental prediction of microbial ASVs.** ASV traits prediction using genetic markers and environmental characters. The numbers denote the average prediction accuracies for microbial ASVs from different treatments across compartments. Only ASVs with heritability ($H^2$) >0.1 were considered in prediction analyses. Boxes span from the first to the third quartiles, centre lines represent the median values and whiskers show data lying within 1.5× interquartile range of the lower and upper quartiles. Data points outside of whiskers represent outliers. CK, control; D, drought; LN, low nitrogen; LP, low phosphorus.

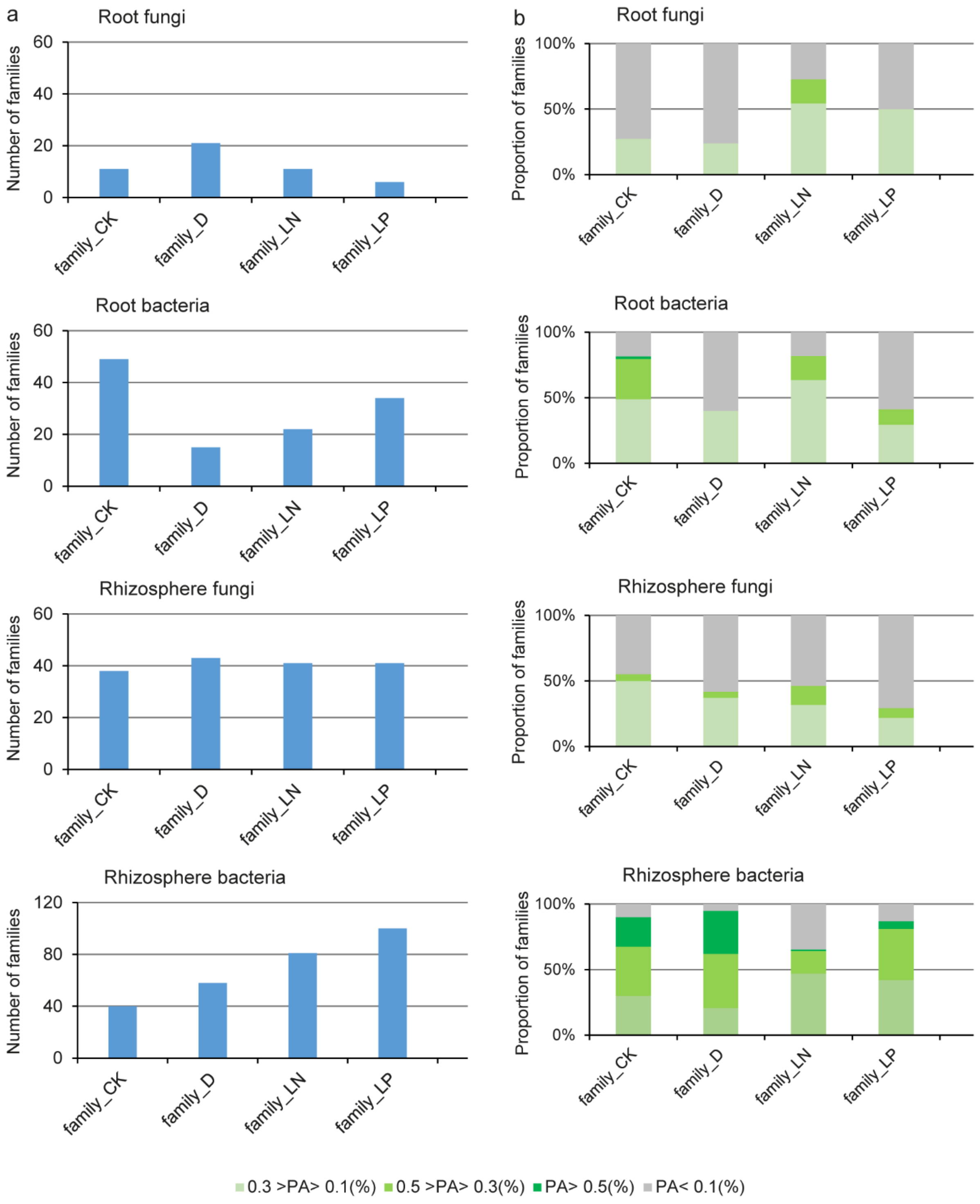


**Supplementary Figure 17. Prediction accuracy of microbial families. a**, Number of highly heritable ($H^2$ >0.1) microbial families. **b**, Degree of prediction accuracies of microbial families from different treatments across compartments. Only families with heritability ($H^2$) >0.1 were considered in prediction analyses. PA, prediction accuracy; CK, control; D, drought; LN, low nitrogen; LP, low phosphorus.

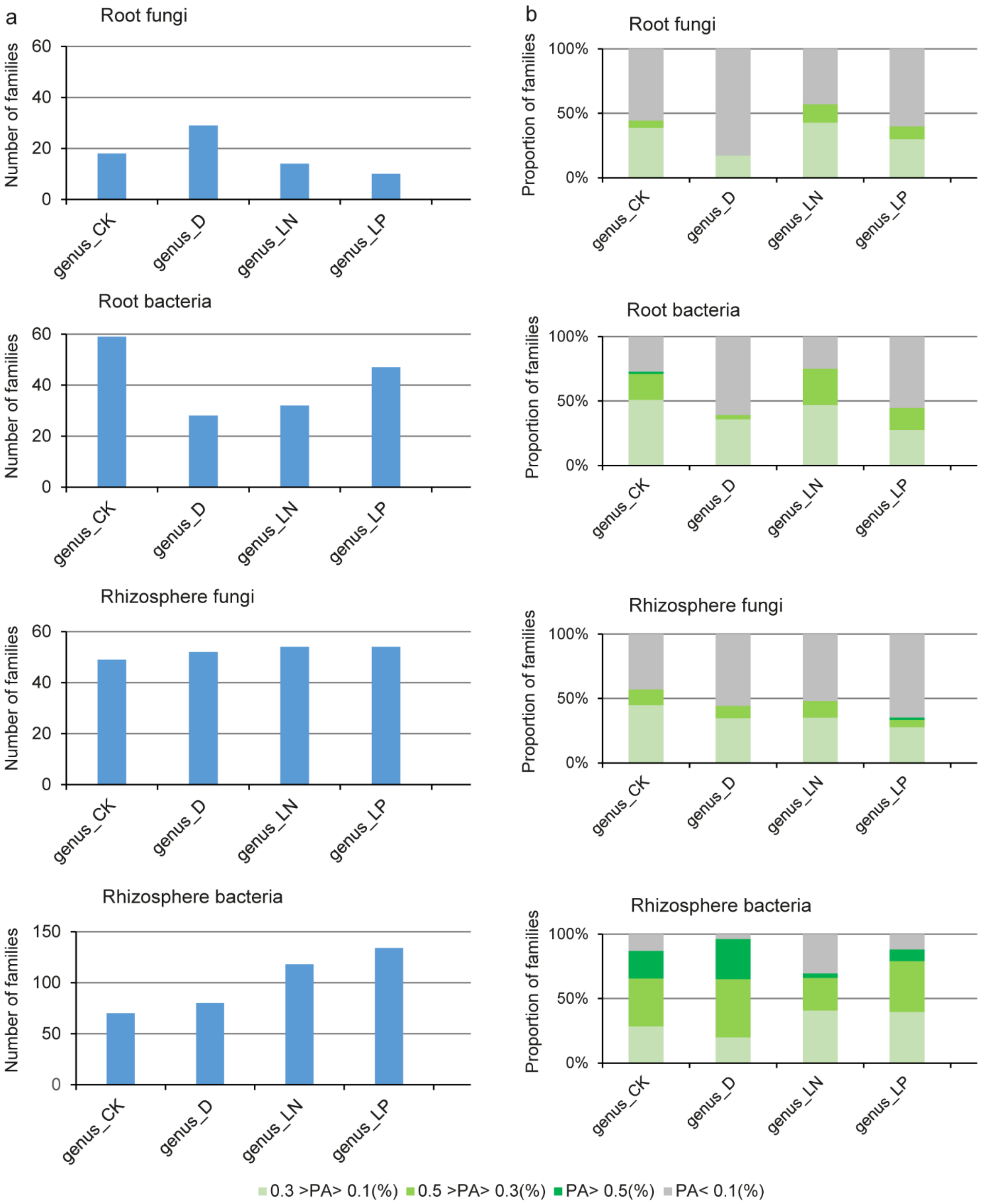


**Supplementary Figure 18. Prediction accuracy of microbial genera. a**, Number of highly heritable ($H^2$ >0.1) microbial genera. **b**, Degree of prediction accuracies of microbial genera from different treatments across compartments. Only genera with heritability ($H^2$) >0.1 were considered in prediction analyses. PA, prediction accuracy; CK, control; D, drought; LN, low nitrogen; LP, low phosphorus.

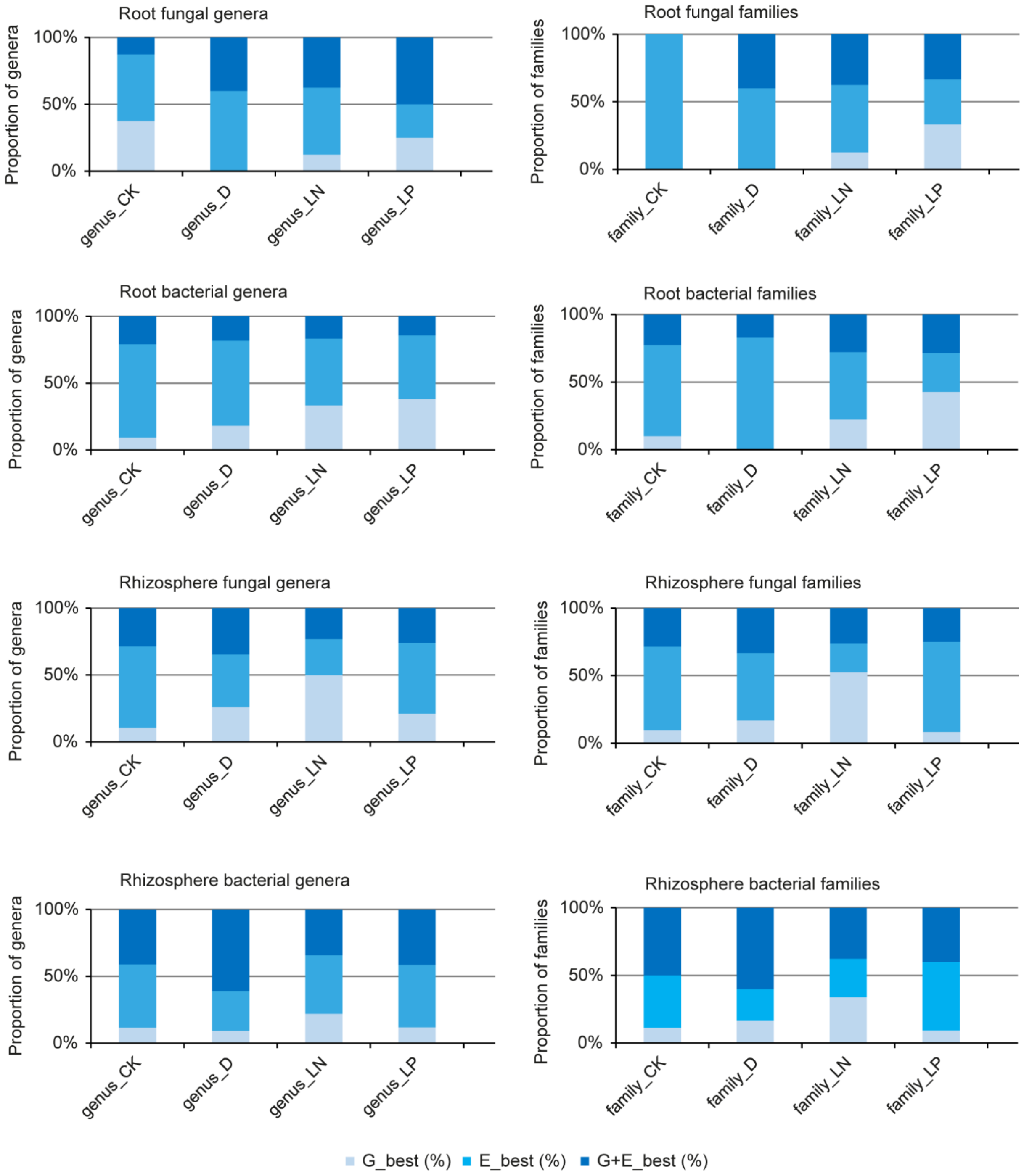


**Supplementary Figure 19. Best prediction patterns of microbial taxa.** G_best indicates best prediction by genetic makers alone. E_best indicates best prediction by environmental characters alone. G+E_best indicates best prediction by combined genetic markers and environmental characters. Only microbial families and genera with heritability ($H^2$) >0.1 were considered in prediction analyses. CK, control; D, drought; LN, low nitrogen; LP, low phosphorus.

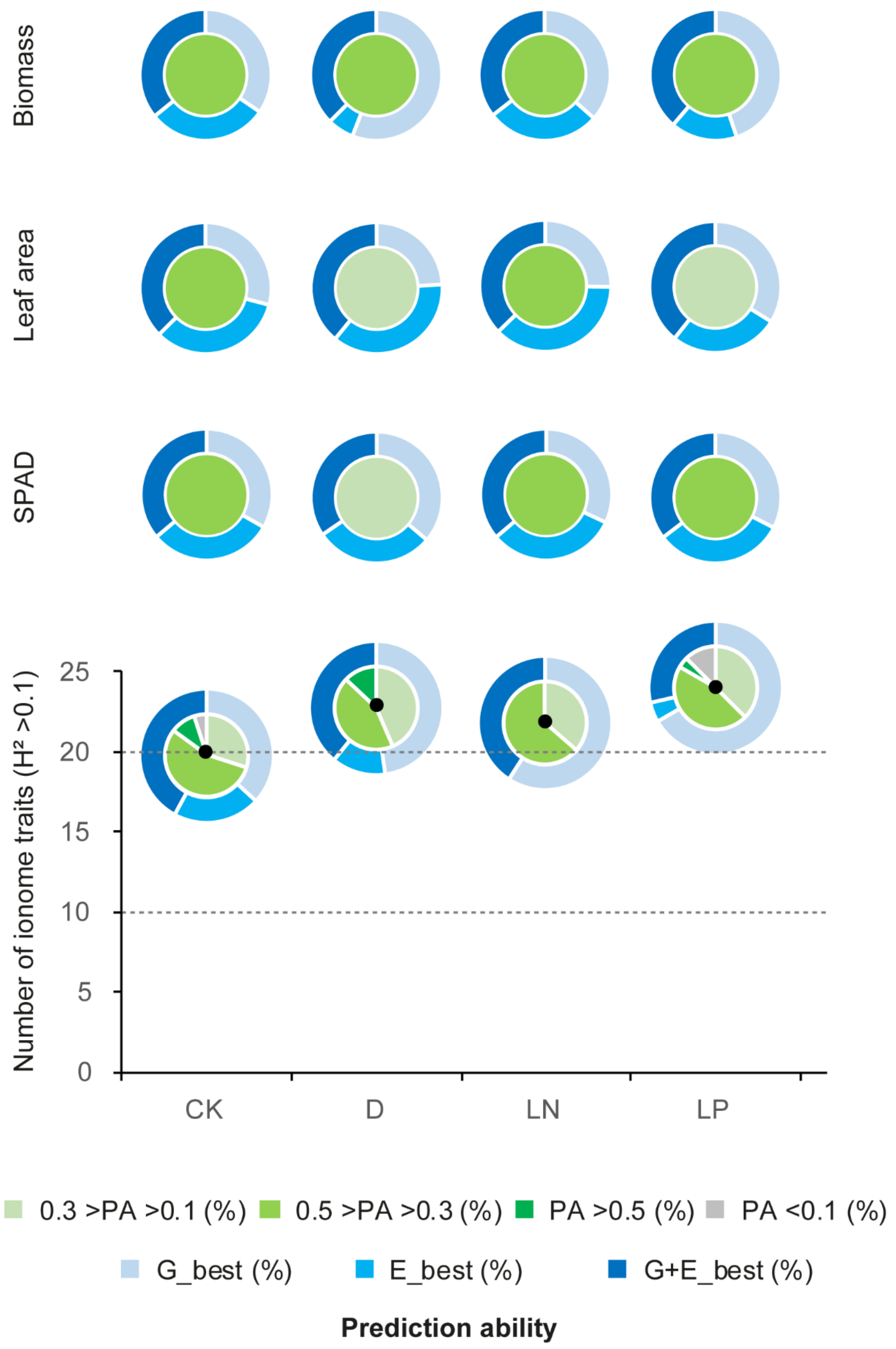


**Supplementary Figure 20. Genomic and environmental prediction of fitness traits.** Fitness traits (biomass, SPAD, leaf area and nutrients) prediction using genetic markers and environmental characters. Inner pie charts describe the degree of prediction accuracies according to proportional traits from different treatments. Outer circles define the best prediction patterns by applying genetic markers (G_best) alone, environmental characters (E_best) alone or a combination of genetic markers and environmental characters (G+E_best). Only traits with heritability ($H^2$) >0.1 were considered in prediction analyses. PA, prediction accuracy. CK, control; D, drought; LN, low nitrogen; LP, low phosphorus.

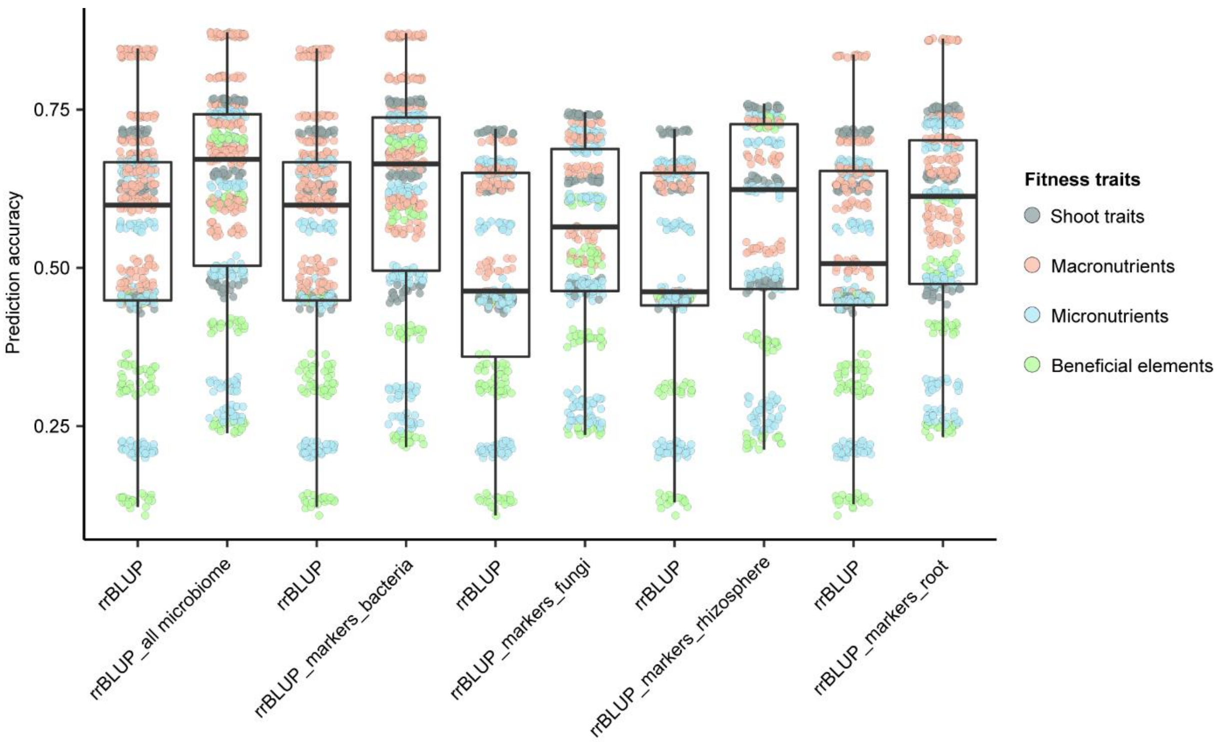


**Supplementary Figure 21. Fitness traits prediction results using host genetics data alone (rrBLUP) and both genetic and microbiome data.** In the figure, different fitness traits were aggregated into one boxplot. In addition to the SNP marker data, the following microbiome data were incorporated into the prediction models: 1) all the microbiomes, including bacteria and fungi from both rhizosphere and root compartments, 2) bacteria from both rhizosphere and root compartments, 3) fungi from both compartments, 4) bacterial and fungi from the rhizosphere, and 5) bacteria and fungi from the root.

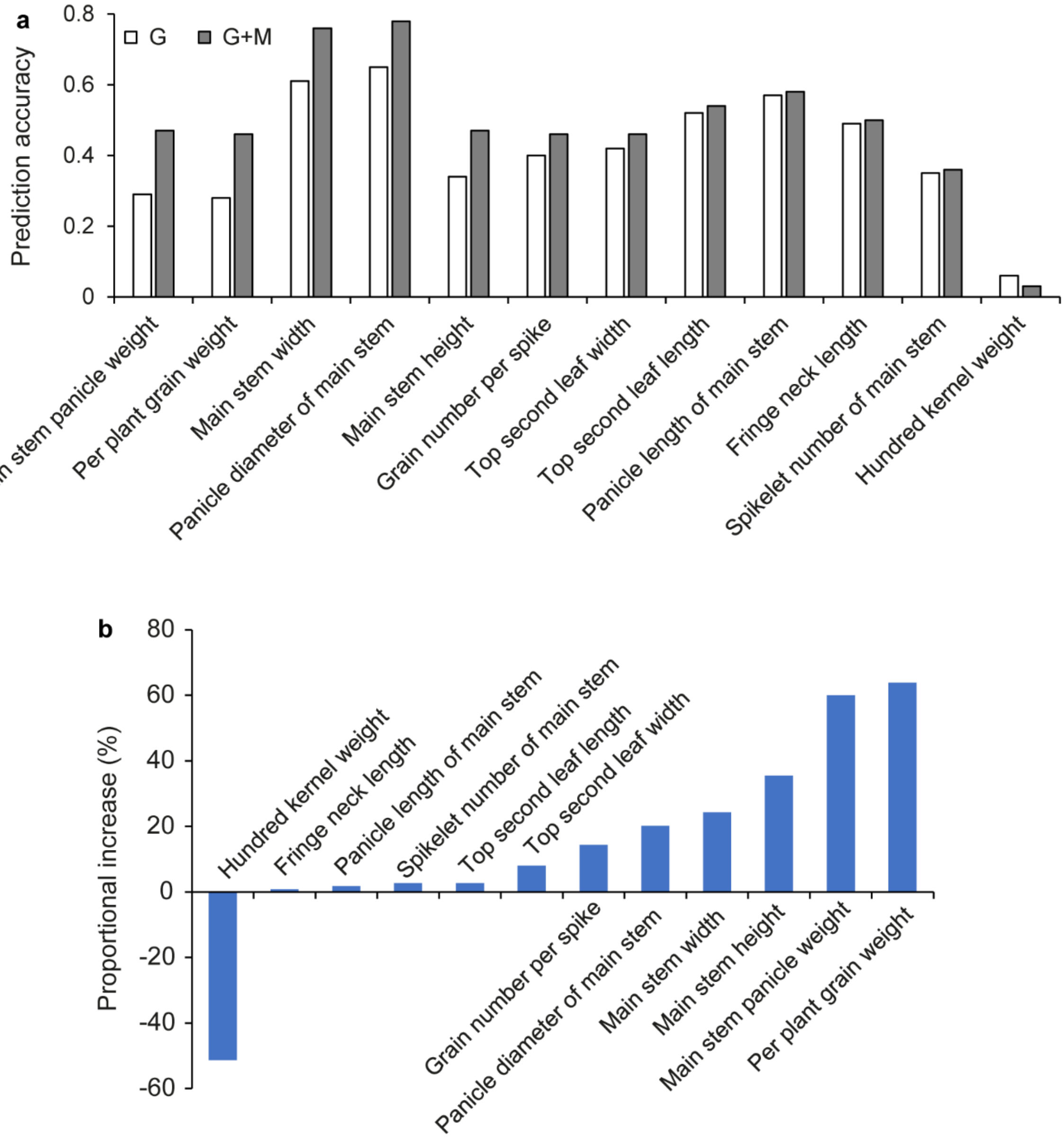


**Supplementary Figure 22. Agronomic traits prediction using genetic markers and microbiome traits in foxtail millet. a**, Prediction accuracy for different agronomic traits using genetic (G) markers alone or combined genetic and microbiome traits (G+M). **b**, Proportional increase of prediction accuracy for different agronomic traits using G+M mode over G mode. 12 agronomic traits were extracted from the publication by Wang et al. 2022 and predicted using genetic markers or combined markers and microbiome traits.

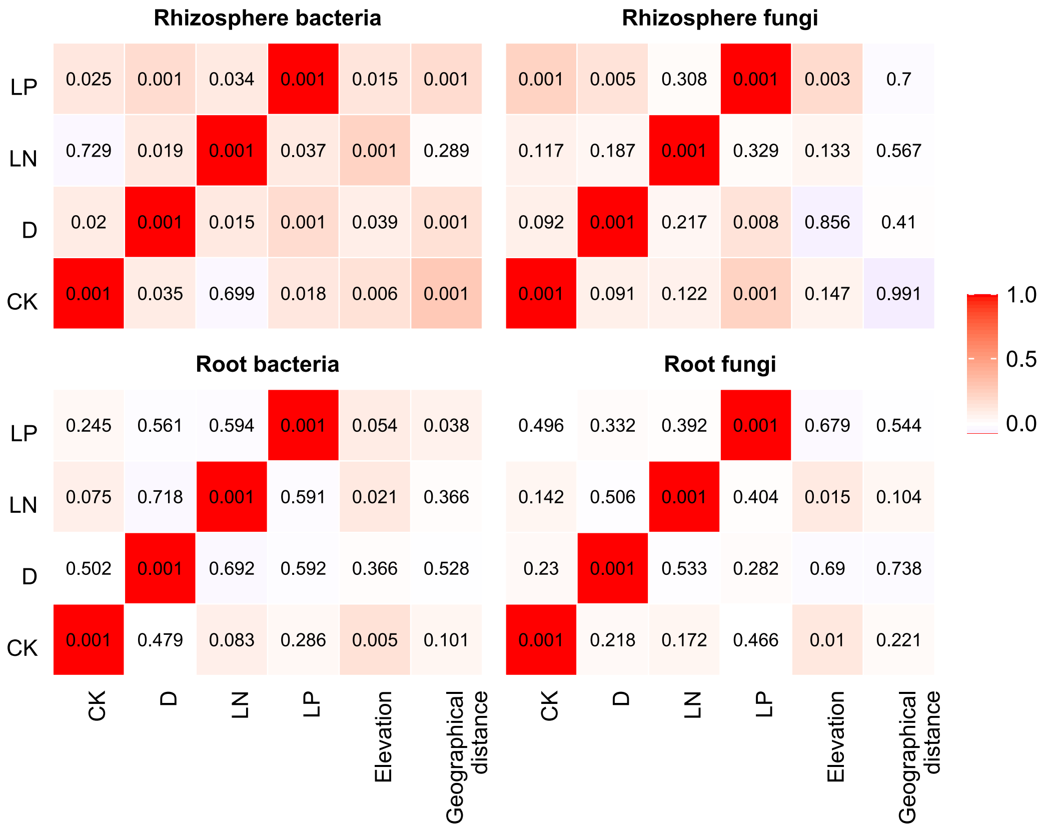


**Supplementary Figure 23. Correlation between microbial communities and local environments.** Mantel's statistical tests were performed between microbiome features and environmental characters (elevation and geographical distance). Significances were indicated by exact $p$ values based on 999 permutations.

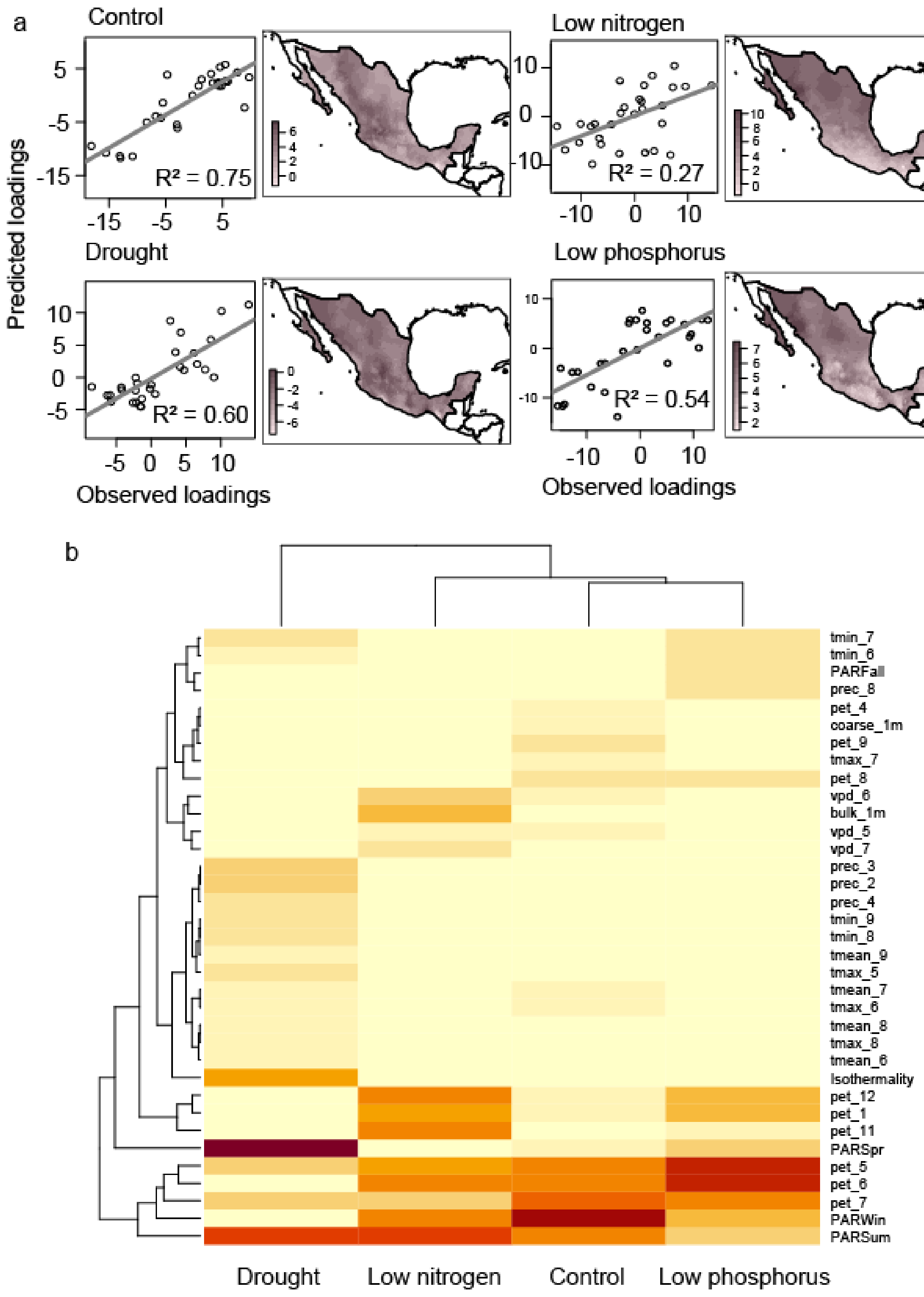


**Supplementary Figure 24. Prediction of rhizosphere bacteria PC2. a,** Random forest approach can well predict the microbiome PC2 in the rhizosphere under different conditions. **b**, Heatmap indicates the importance of environmental variables in PC2 predictions across treatments.

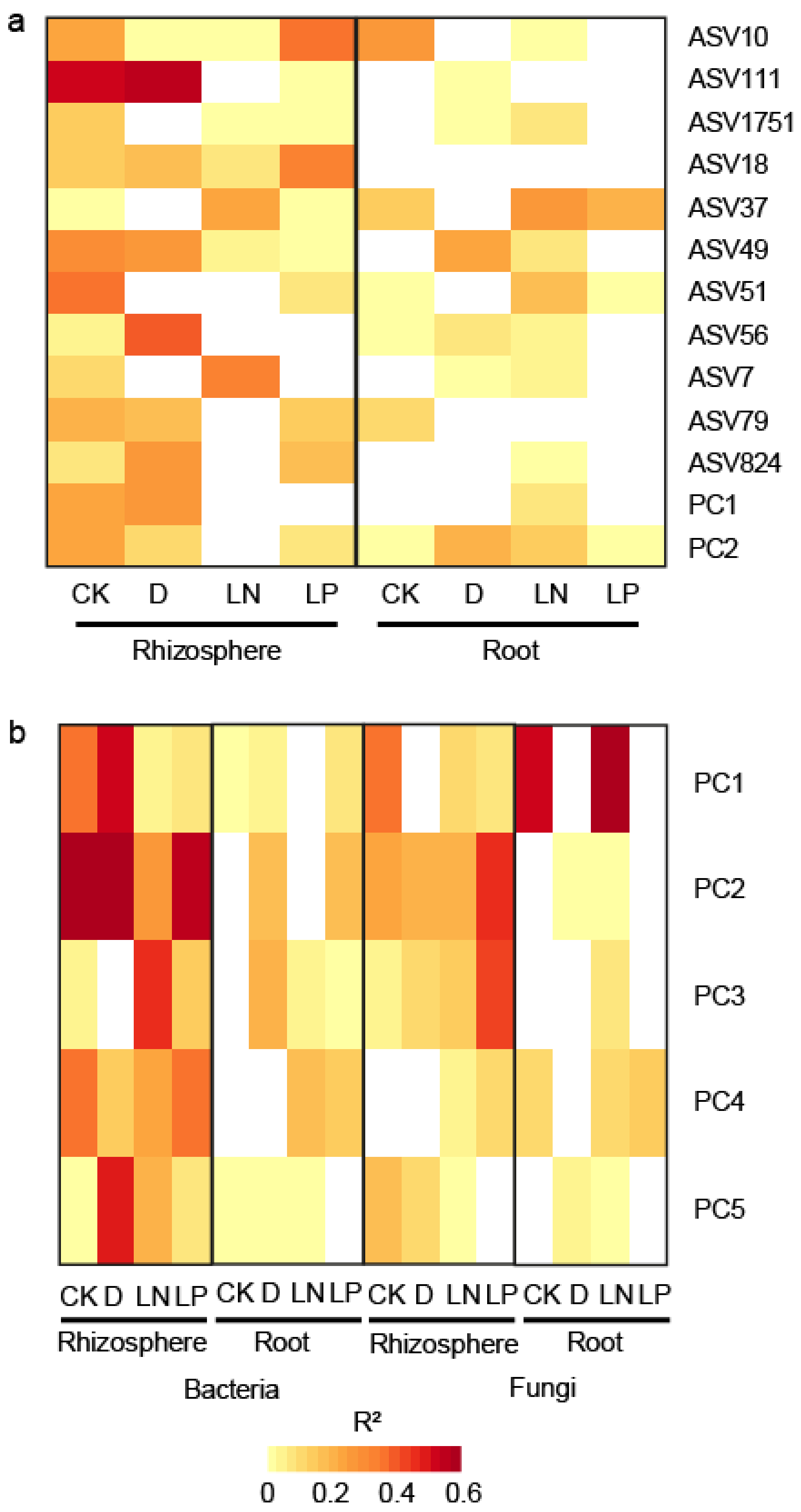


190 **Supplementary Figure 25. Microbiome prediction. a**, Prediction of microbial abundance by using
191 individual *Massilia* ASVs. **b**, Prediction of microbial abundance by using the whole microbiome (top 5
192 PCs).

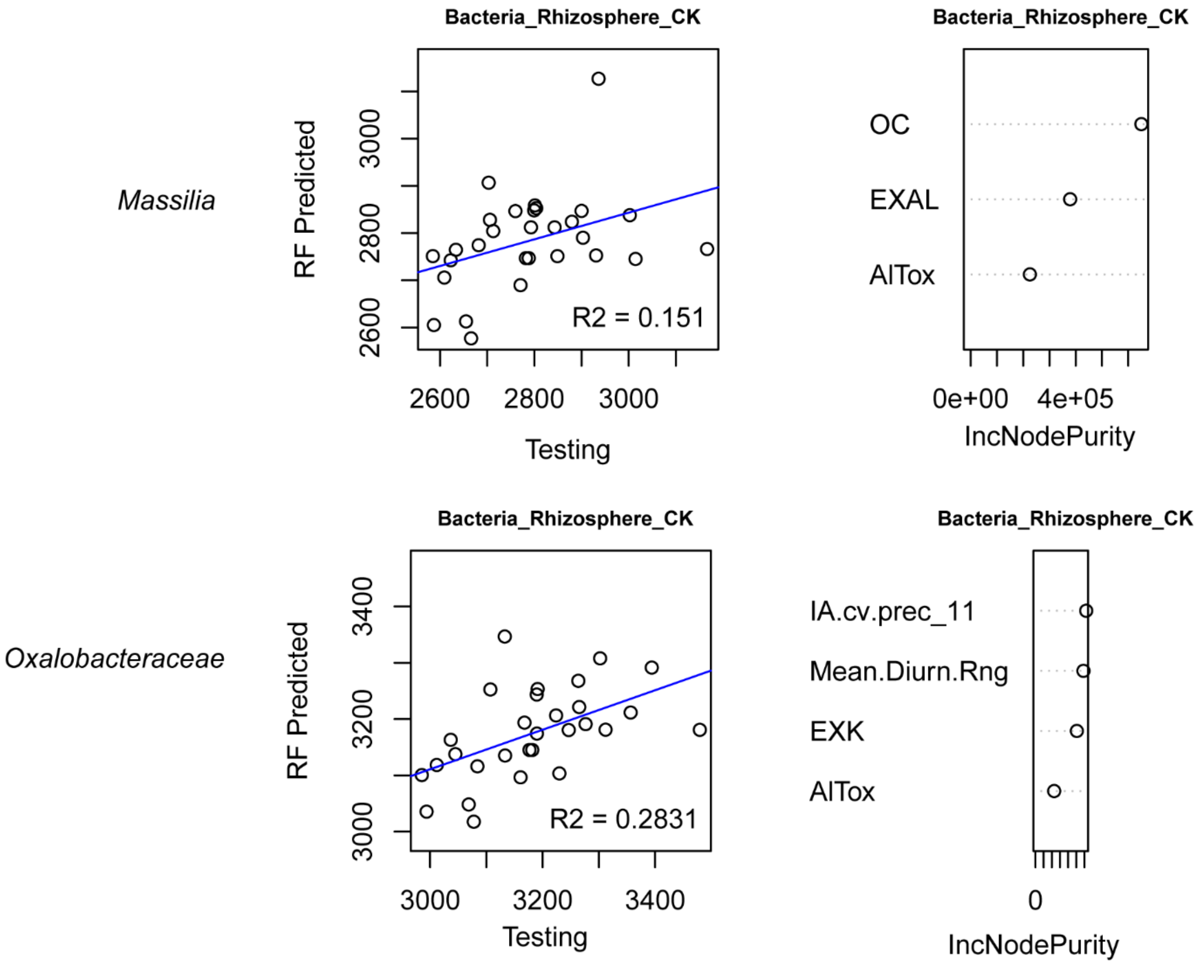


**Supplementary Figure 26. Random forest modelling correlation of testing taxa.** The environmental characters can be referred to from Supplementary Dataset 12.

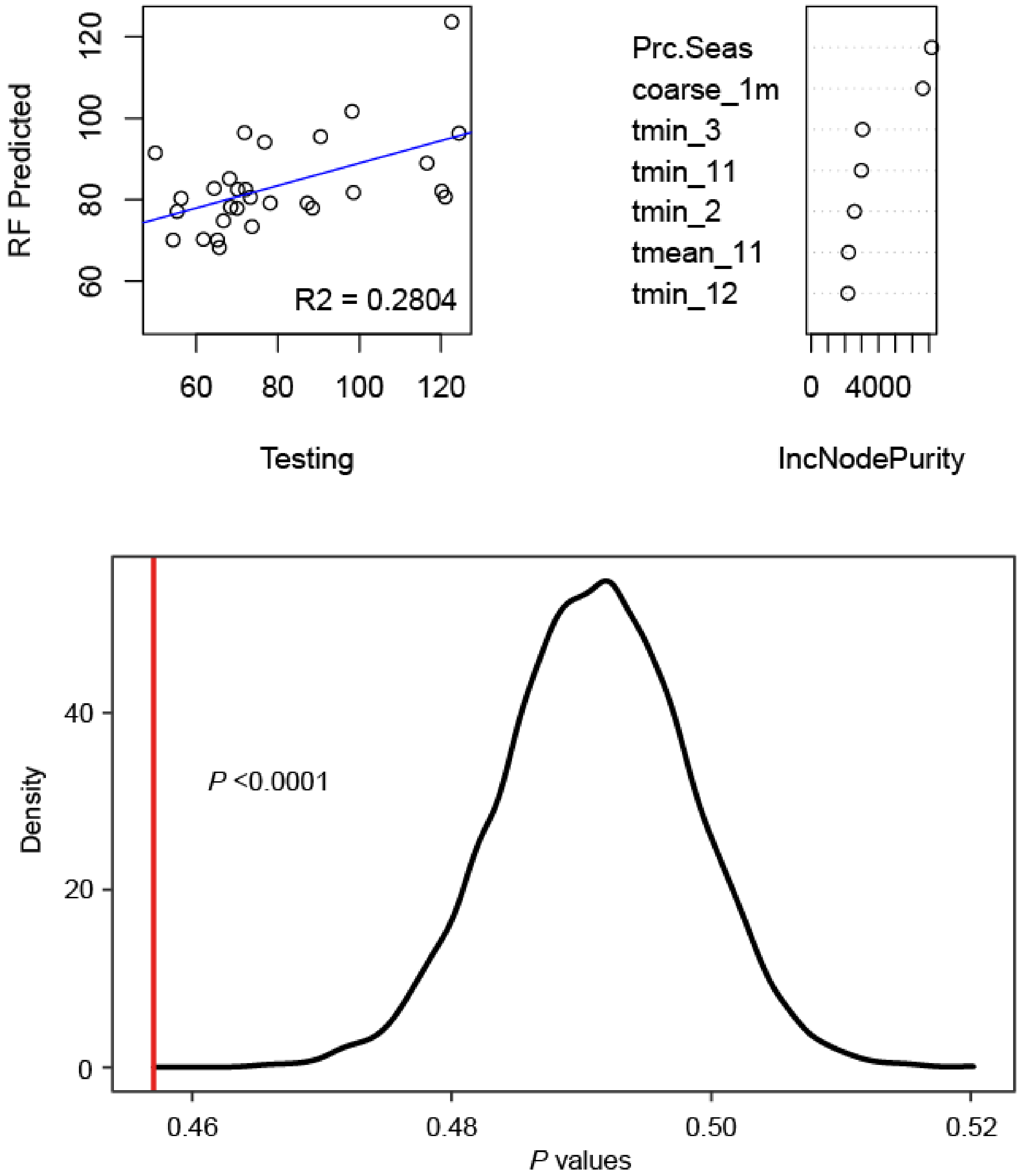


**Supplementary Figure 27. Random forest modelling correlation of ASV37 in root under nitrogen-poor conditions.** The environmental characters can be referred to from Supplementary Dataset 12. Permutation test results showing significantly lower median *p*-value for SNPs from the phenotypic GWAS that are around 200kb of the top 100 SNPs of predicted GWAS (red vertical line) than the median p-value of random selected SNPs of phenotypic GWAS, based on 10000 permutations.

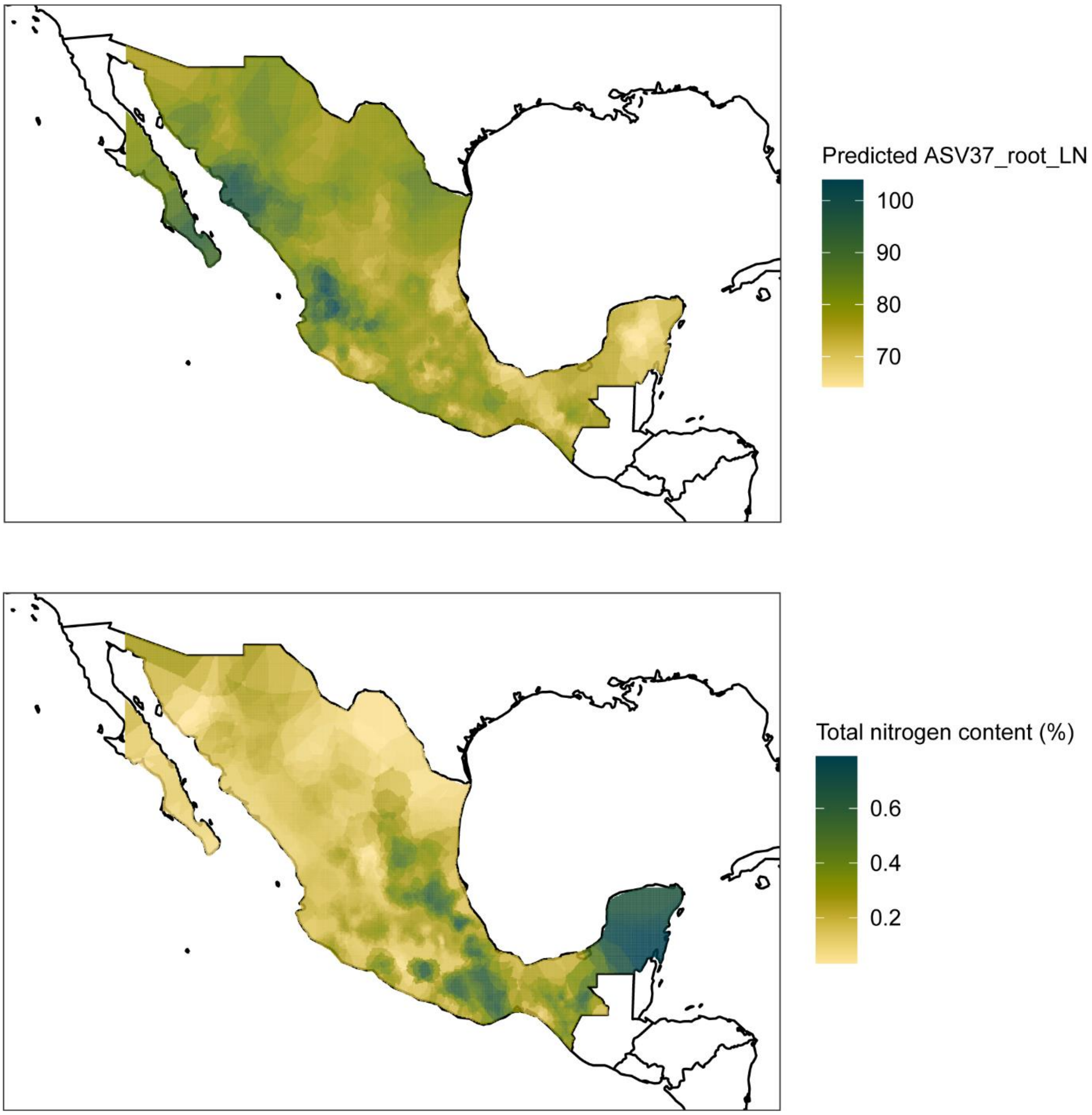


**Supplementary Figure 28. Maps showing the spatial variation of targeted ASV and total soil nitrogen contents based on 1781 landraces.** Top panel: Predicted relative abundance of ASV37 in roots under low nitrogen; Bottom panel: Distribution of total nitrogen content in soils. Green shades correspond to higher values, whereas yellow shades correspond to lower values.

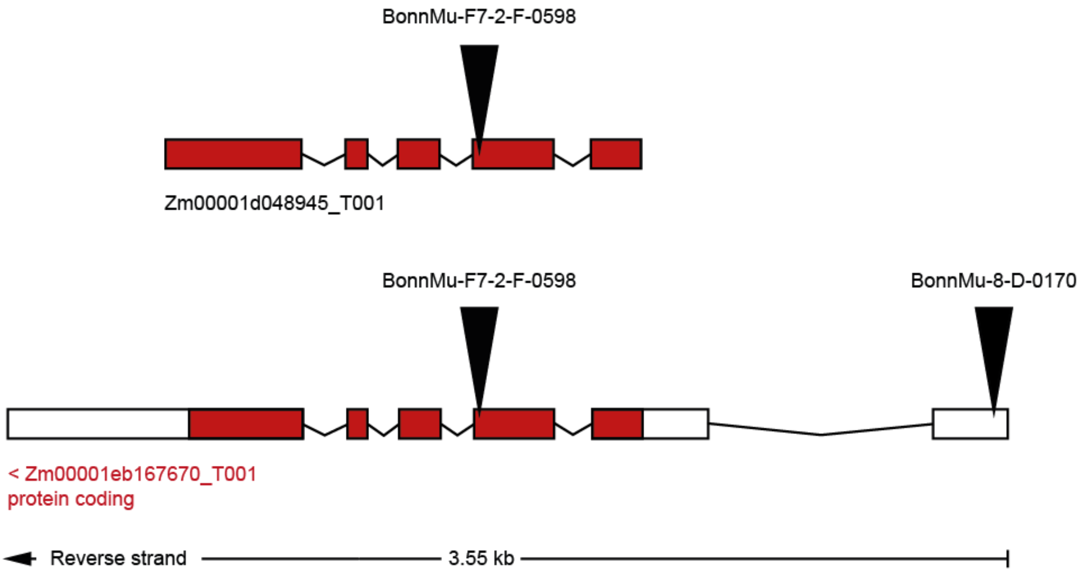


**Supplementary Figure 29. Transposon-tagged mutations of gene Zm00001d048945 (version 4).** Induced maize mutants of the BonnMu resource derive from Mutator-tagged $F_2$-families in the genetic backgrounds B73 and F7. We identified two insertion lines, BonnMu-8-D-0170 (B73) and BonnMu-F7-2-F-0598 (F7), harboring insertions 1,264 bp upstream of the start codon ATG and in the second exon of Zm00001eb167670 (version 5), respectively.

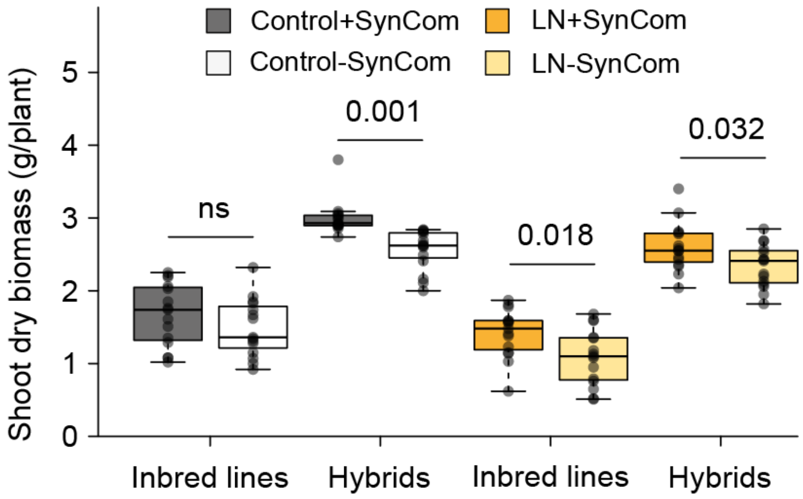


**Supplementary Figure 30. Shoot growth in the presence/absence of an *Oxalobacteraceae*-based synthetic community of inbred lines and hybrids under different soil nitrogen conditions.** Exact *p* values were calculated using two-tailed unpaired *t*-tests according to *F*-test results. Boxes span from the first to the third quartiles, centre lines represent the median values and whiskers show data lying within 1.5× interquartile range of the lower and upper quartiles. Data points at the ends of whiskers represent outliers.

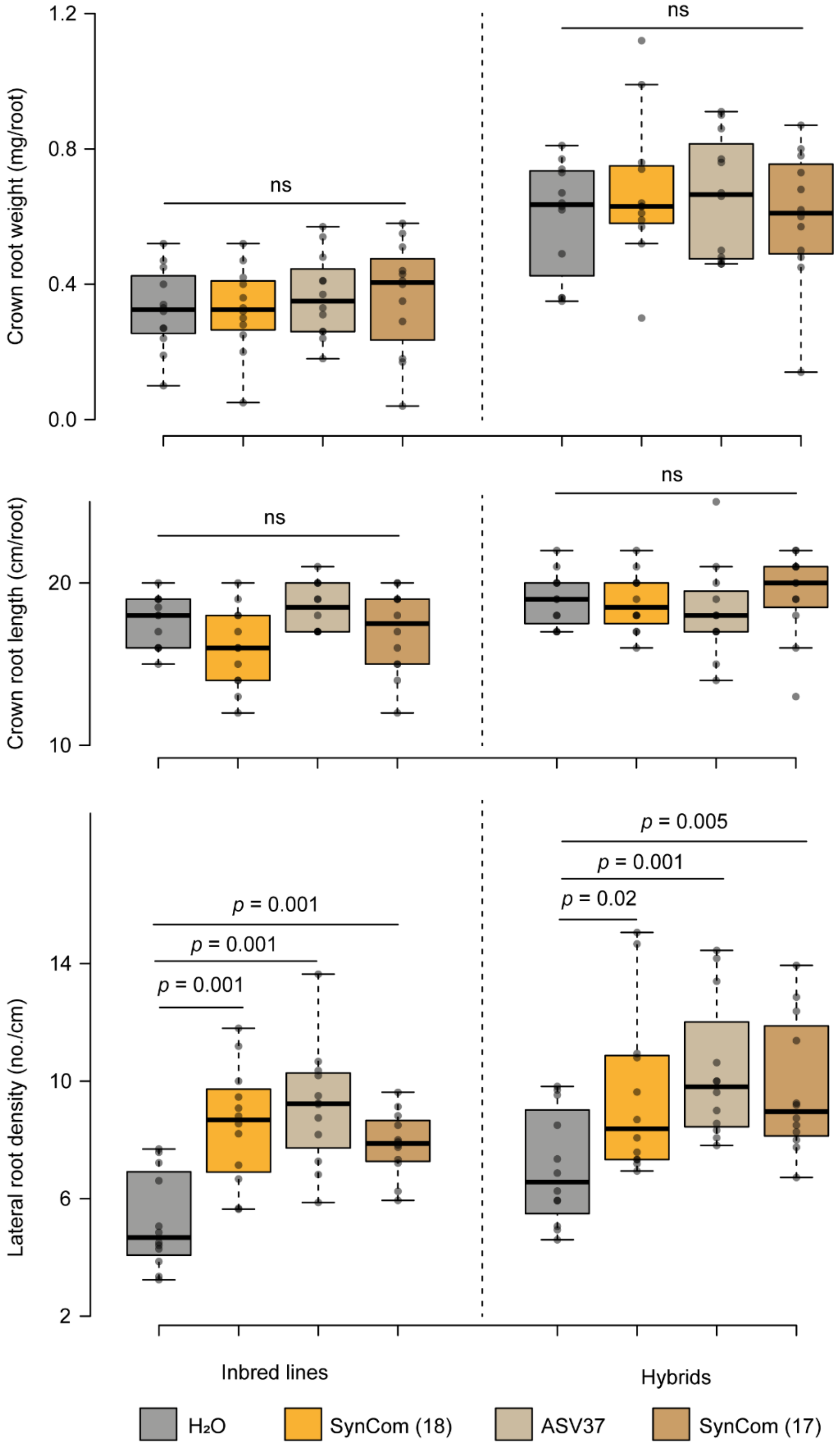


**Supplementary Figure 31. Root traits in the presence/absence of *Oxalobacteraceae* of inbred lines and hybrids under nitrogen-poor conditions.** Exact *p* values were calculated using one-way ANOVA by turkey HSD. ns, not significant. Boxes span from the first to the third quartiles, centre lines represent the median values and whiskers show data lying within 1.5× interquartile range of the lower and upper quartiles. Data points at the ends of whiskers represent outliers.

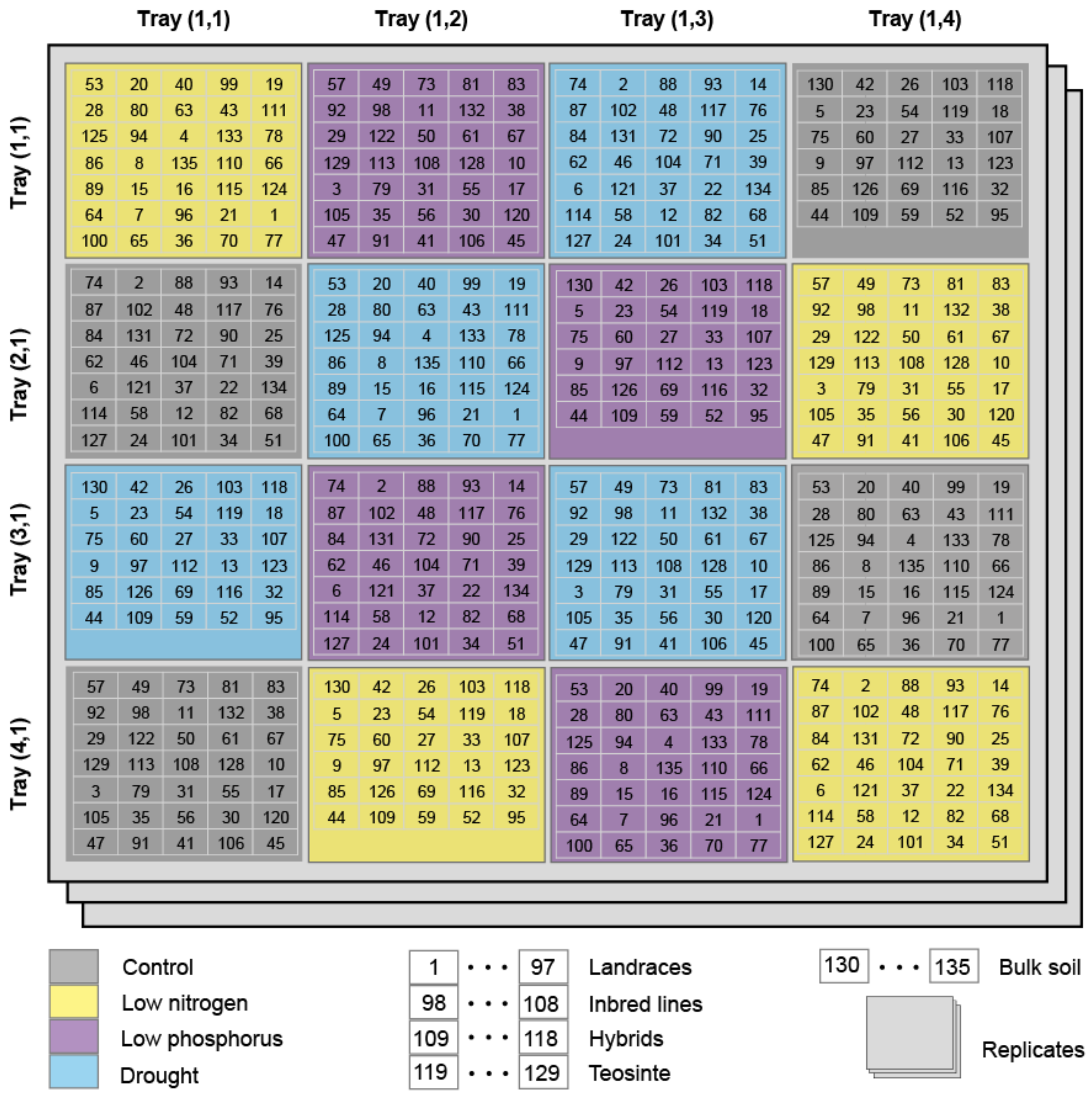


**Supplementary Figure 32. Experimental design.** The experiment was laid out in a split plot design comprising four treatments: (1) fully fertilized control soil, (2) low nitrogen (LN) soil with no nitrogen fertilizer applied, (3) low phosphate (LP) soil with no phosphate fertilizer applied and (4) control soil with drought (D) treatment. Different numbers were allocated to the different genotypes: landraces (1-97), inbred lines (98-108), hybrids (109-118) and teosinte (119-129). Six pots (130-135) without plants were set up as ‘bulk soil’ samples. Each tray comprises 35 pots with different germplasm groups and bulk soil pots. The pot distribution in each tray was randomized. The whole set-up was repeated three times as three biological replicates.

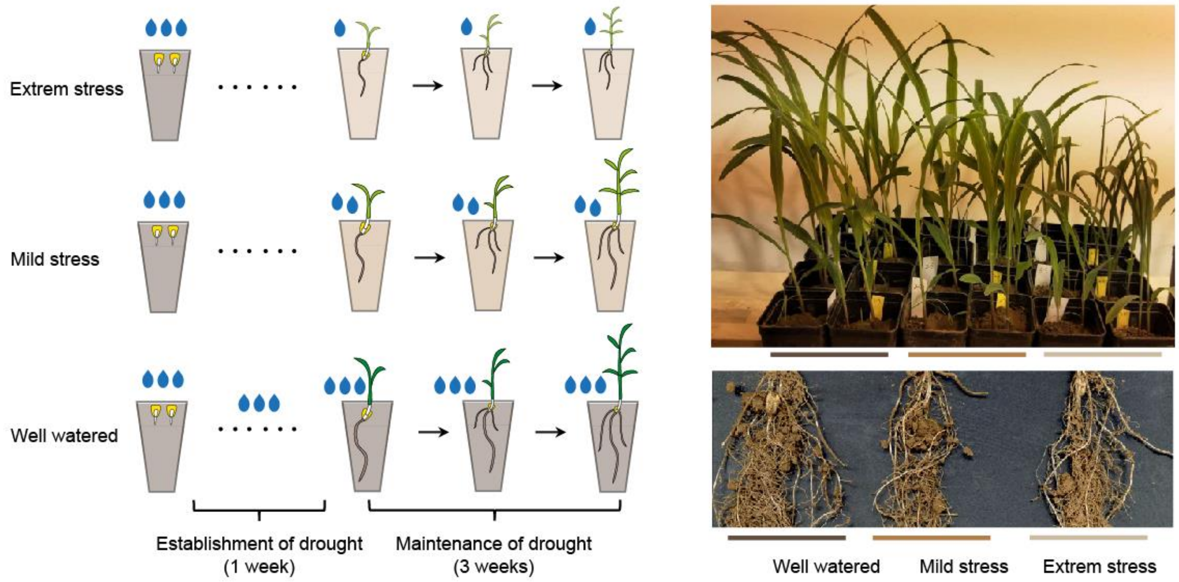


**Supplementary Figure 33. Drought preliminary experiment.** The left panel indicates the three drought experiments with different water levels. The right panel indicates the shoot and root phenotypes grown under three different drought levels.

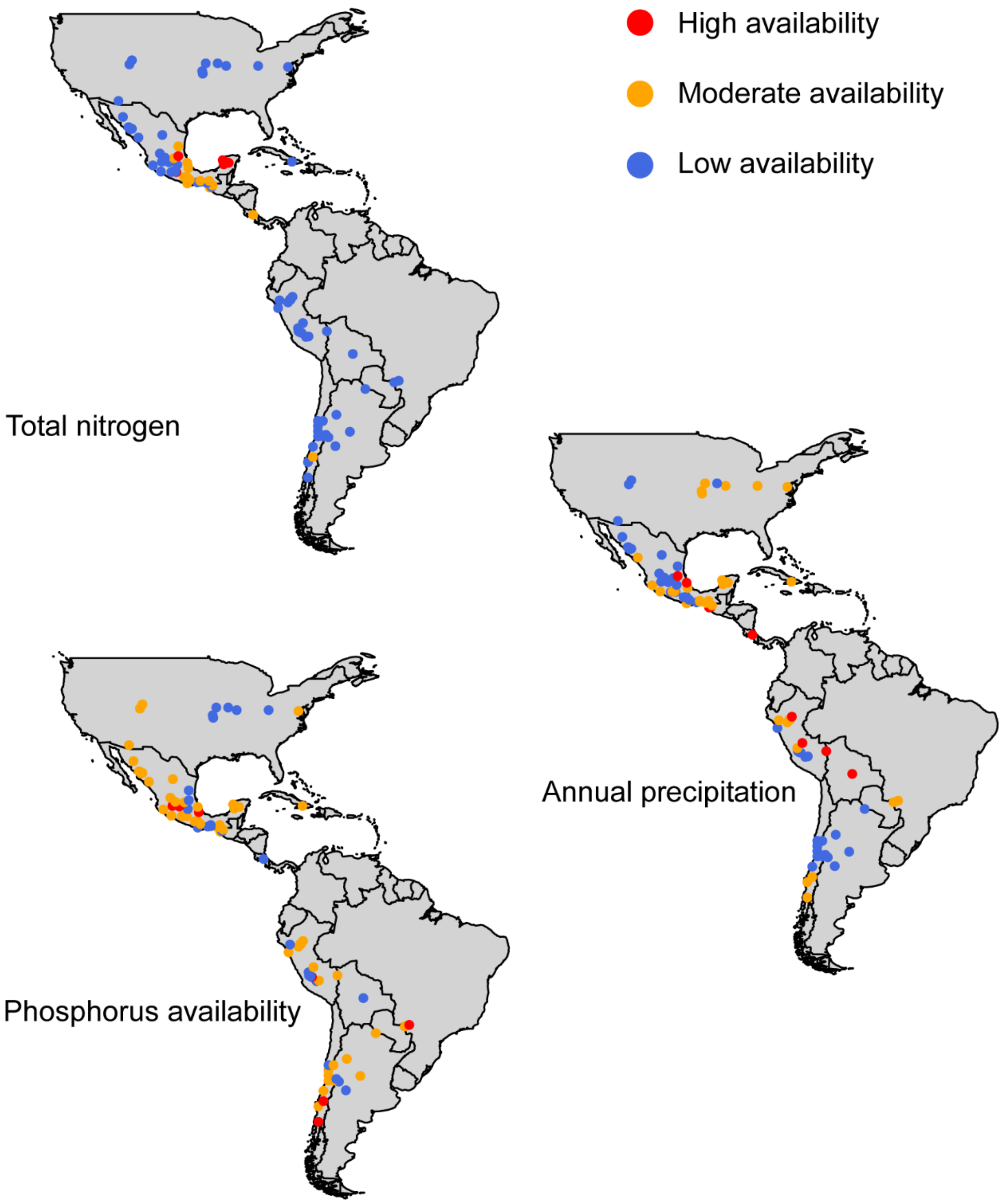


**Supplementary Figure 34. Availability of nutrients and precipitation for landraces collected in their natural habitats.** The original data is provided in Supplementary Dataset 12.

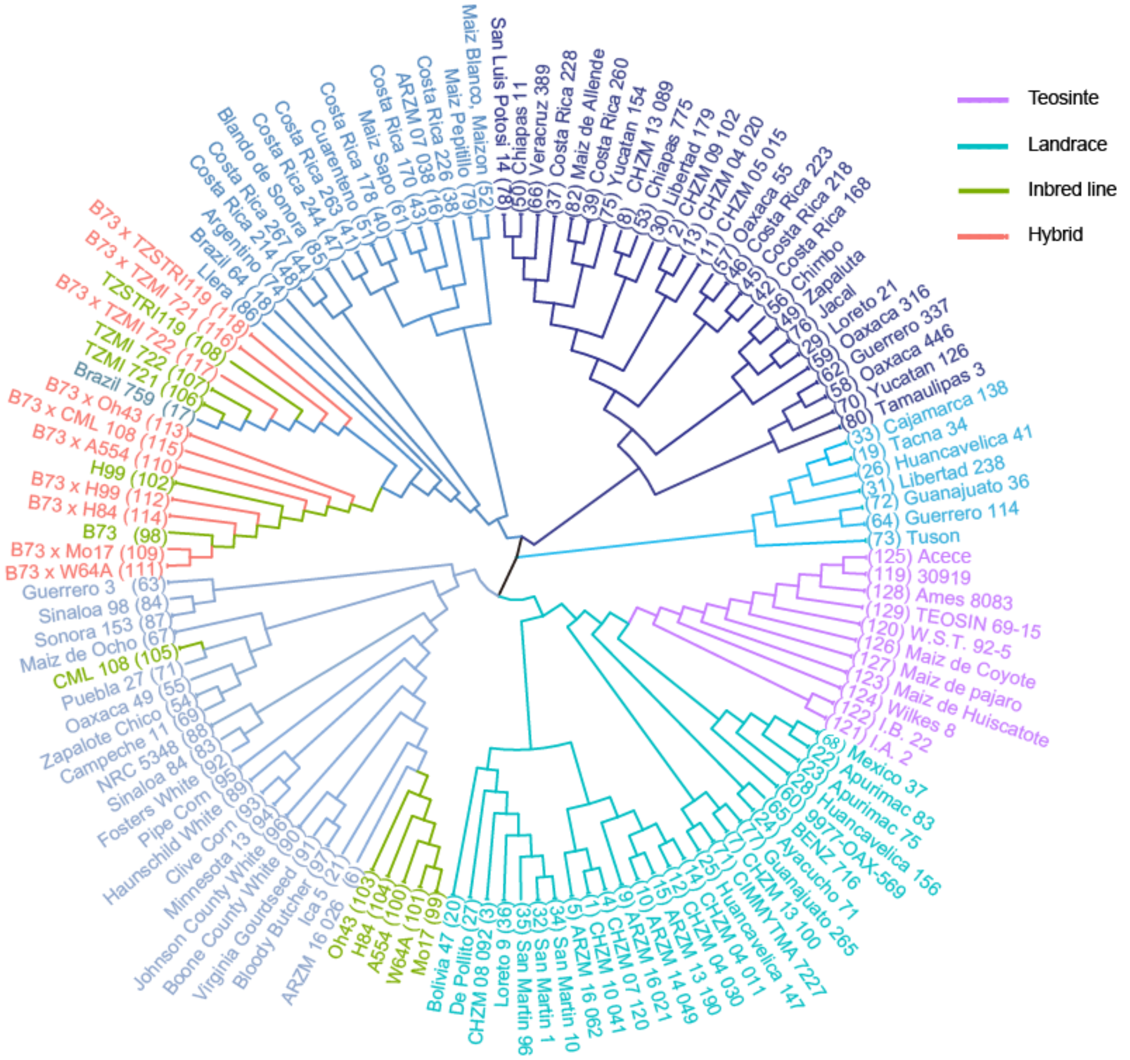


243

244 **Supplementary Figure 35. Phylogenetic relationship of the 129 maize genotypes used in this**
245 **study.** A neighbour-joining tree was produced using SNP markers. Different colours correspond to
246 different germplasm groups. Different shades of blue represent different sources of local landraces.

**Supplementary Table 1 Basic soil physical and chemical properties.**

The values presented are mean ± standard error of all samples for three nutrient treatments collected from Dikopshof in 2019. Means followed by the same letter are not significantly different at $p$ <0.05 according to LSD (ANOVA, turkey HSD, $n$ = 5).

| Soil properties | Control | Low nitrogen | Low phosphorus |
|---|---|---|---|
| Total N (%) | 0.094 ± 0.0027a | 0.076 ± 0.00088b | 0.087 ± 0.00053a |
| Conc. P (mg/kg) | 69.23 ± 1.61a | 67.55 ± 1.06a | 24.50 ± 1.39b |
| Conc. K (mg/100g ) | 10.43 ± 0.093b | 10.97 ± 0.064a | 8.13 ± 0.072c |
| Conc. Fe (mg/100ml) | 35.52 ± 0.62a | 38.05 ± 0.53a | 30.44 ± 0.74b |
| Conc. Mn (mg/100ml) | 2.42 ± 0.037a | 2.49 ± 0.20a | 2.05 ± 0.036a |
| Conc. Cu (mg/100ml) | 0.035 ± 0.00066a | 0.035 ± 0.00078a | 0.035 ± 0.00090a |
| Conc. Zn (mg/100ml) | 0.071 ± 0.00058ab | 0.080 ± 0.0014a | 0.069 ± 0.0034b |
| pH | 6.74 ± 0.08b | 7.23 ± 0.07a | 6.97 ± 0.07ab |
| C (%) | 0.81 ± 0.014a | 0.71 ± 0.0062b | 0.83 ± 0.032a |
| Total organic carbon (%) | 0.60 ± 0.062a | 0.56 ± 0.043a | 0.69 ± 0.010a |
